# Next-generation interaction proteomics for quantitative Jumbophage-bacteria interaction mapping

**DOI:** 10.1101/2023.01.13.523954

**Authors:** Andrea Fossati, Deepto Mozumdar, Claire Kokontis, Melissa Mèndez-Moran, Eliza Nieweglowska, Adrian Pelin, Yuping Li, Baron Guo, Nevan J. Krogan, David A. Agard, Joseph Bondy-Denomy, Danielle L. Swaney

**Author notes:** Contributing authors. These authors contributed equally to this work (Andrea Fossati, Deepto Mozumdar).

## Abstract

Host-pathogen interactions (HPIs) are pivotal in regulating establishment, progression, and outcome of an infection. Affinity-purification mass spectrometry has become instrumental for the characterization of HPIs, however the targeted nature of exogenously expressing individual viral proteins has limited its utility to the analysis of relatively small pathogens. Here we present the use of co-fractionation mass spectrometry (SEC-MS) for the high-throughput analysis of HPIs from native viral infections of two jumbophages (*ϕ*KZ and *ϕ*PA3) in *Pseudomonas aeruginosa*. This enabled the detection *>*6000 unique host-pathogen and *>*200 pathogen-pathogen interactions for each phage, encompassing *>*50% of the phage proteome. Interactome-wide comparison across phages showed similar perturbed protein interactions suggesting fundamentally conserved mechanisms of phage predation within the KZ-like phage family. Prediction of novel ORFs revealed a *ϕ*PA3 complex showing strong structural and sequence similarity to *ϕ*KZ nvRNAp, suggesting *ϕ*PA3 also possesses two RNA polymerases acting at different stages of the infection cycle. We further expanded our understanding on the molecular organization of the virion packaged and injected proteome by identifying 23 novel virion components and 5 novel injected proteins, as well as providing the first evidence for interactions between KZ-like phage proteins and the host ribosome. To enable accessibility to this data, we developed PhageMAP, an online resource for network query, visualization, and interaction prediction (https://phagemap.ucsf.edu/). We anticipate this study will lay the foundation for the application of co-fractionation mass spectrometry for the scalable profiling of hostpathogen interactomes and protein complex dynamics upon infection.

## Introduction

Protein-protein interactions (PPIs) are the fundamental building blocks of cellular complexity and their perturbation and rewiring have profound effects on the proteome and cell fate. During an infection, the interactions between host and pathogen proteome are pivotal in regulating pathogen tropism, infection progression and, ultimately, infection outcome. Host-pathogen interaction (HPI) mapping using affinity-purification mass spectrometry (AP-MS) has been instrumental in identifying host targeted processes[1–5] and, recently, to predict potential therapeutic targets during the SARS-CoV-2 pandemic[6–8].

Despite the successes of AP-MS for mapping HPIs, the exogenous expression and purification of individual pathogen proteins limits our ability to characterize HPIs under native expression levels, and quantify how these interactions are regulated in the context of the full pathogen protein repertoire during an infection. The targeted nature of AP-MS also precludes the detection of downstream rearrangements in protein complexes beyond the viral protein of interest. Lastly, AP-MS is a labor-intensive process that requires the generation of numerous plasmids and hundreds or thousands of individual purifications to comprehensively probe protein-protein interactions for an entire viral proteome. This limits the scalability of AP-MS for the characterization HPIs for larger viruses or bacteria which express hundreds or thousands of proteins.

As a result, small eukaryotic viruses have been prioritized in host-pathogen interaction studies[6, 9], thus extensive knowledge on interactions between larger prokaryotic viruses (bacteriophages) and their host is currently missing. This class of bacterial viruses hold great potential for treatment of multi-drug resistant bacteria which have increasingly been reported in the last two decades[10]. However, without a thorough understanding of putative interactions and functions of the phage gene products, it will be challenging to inform the rational design of the next generation of phage therapeutics.

To bridge this gap, here we have applied co-fractionation mass spectrometry using size-exclusion chromatography, coupled with fast data-independent acquisition MS (SEC DIA-MS),[11] to generate two phage-bacteria interactomes and to measure host PPI rewiring upon phage infection in *Pseudomonas aeruginosa*. Specifically, we provide the first interactome of two KZ-like phages (*ϕ*KZ[12] and *ϕ*PA3[13]) which are archetype Jumbophages that possess large genomes (*>*300 genes), with very limited organization of genes by function, hence lacking synteny. Unique to this family of phages is the presence of a large proteinaceous shell acting analogous to the eukaryotic nucleus thus decoupling transcription from translation. This structure confers resistance to several bacterial antiphage systems such as CRISPR[14, 15] and has a fundamental role in infection establishment[16] and virion production[17]. Through the prediction of PPIs using deep learning and structural modeling, we derived system-level maps of Jumbophage infection encompassing a large fraction of the phage and bacterial host proteome. These host-pathogen interaction maps substantially extend previous knowledge on Jumbophage predation and provide the first application of co-fractionation mass spectrometry for host-pathogen interaction profiling.

## Results

### A cross-phage study of viral infection cycle

To understands HPIs that mediate phage infection, we infected *Pseudomonas aeruginosa* (strain PAO1) with either the *ϕ*KZ or *ϕ*PA3 bacteriophage for 60 minutes in biological duplicate. To control for virion protein complexes (i.e complexes present within the phage itself), parallel experiments were also performed using a mutant PAO1 strain that emerged under phage selection (KZ resistant mutant)[18] that resists infection (Fig.1A), dubbed ‘PAO1 control’. This strain expresses significantly lower levels of FliC protein (the major structural unit of the flagellum), a known receptor for *ϕ*KZ[18].

**Fig. 1.**
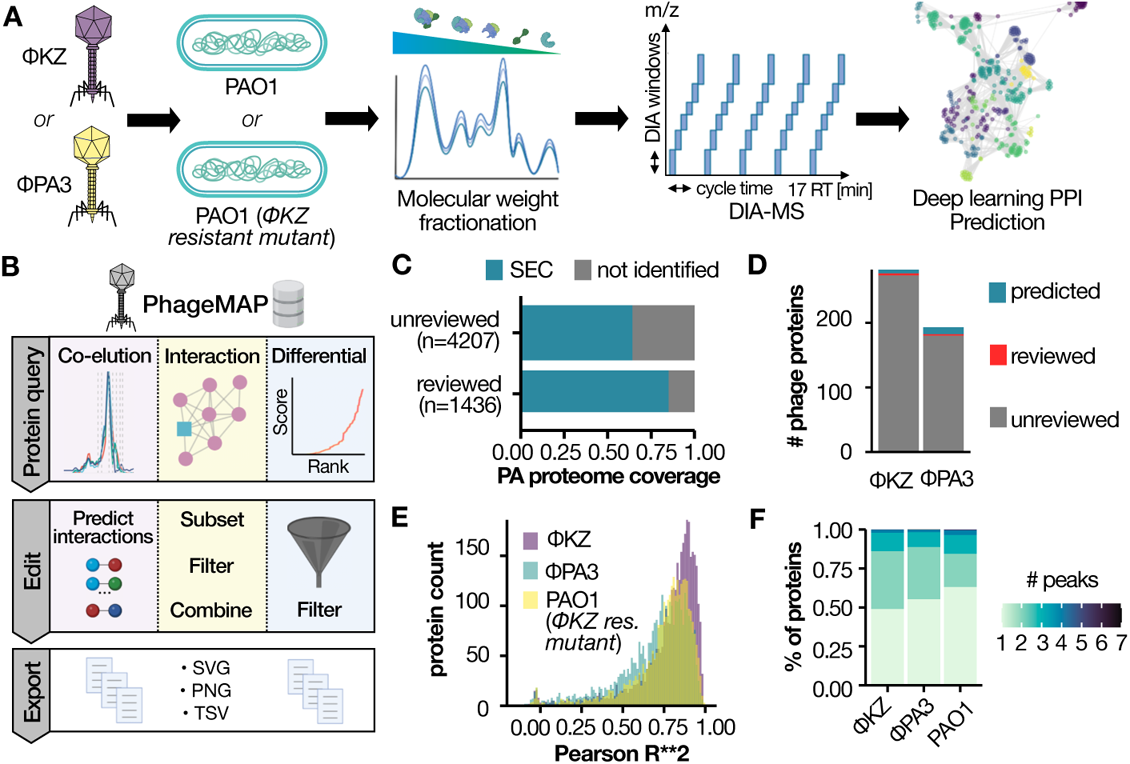
High-throughput interaction proteomics for deep host-pathogen interaction mapping. **A**. SEC-MS workflow and experimental design. **B**. Overview of PhageMAP analysis and workflows. **C**. Recovery of *Pseudomonas* proteome by SEC-MS. **D**. Barplot representing the number of phage proteins identified. **E**. Correlation for all proteins identified in the experiment (n=4132) between the two replicates. **F**. Fractional distribution of the number of SEC peaks across the various phages and host.

Infected cell lysates were fractionated by size-exclusion chromatography, and each fraction (n=72) was analyzed using data independent acquisition MS (DIA-MS) coupled to high-throughput liquid chromatography[19]. To predict host-pathogen interactions, we used a modified version of the PCprophet toolkit[11], where the random forest classifier was replaced with a deep neural network that was trained for PPI prediction using *>*10 million interactions from various co-fractionation experiments[20].

Derived host-pathogen interaction networks have been organized into a user-friendly website, PhageMAP, where users can query proteins of interest to visualize coelution patterns, interactomes, investigate different assembly states of the PAO1 proteome upon phage infection, and export their findings as publication-quality networks or coelution plots (Fig.1B).

This experimental workflow resulted in the high-throughput and comprehensive coverage of both the bacterial and the phage proteomes. Specifically, we detected 3782 PAO1 proteins, covering 83% of the validated SwissProt entries (i.e proteins for which experimental evidence of their existence is available) for the *Pseudomonas* pan-proteome, and 67% of the unreviewed entries (Fig.1B). Likewise, we detected 280 proteins for *ϕ*KZ and 198 proteins for *ϕ*PA3, covering 75% and 53% of their proteomes, respectively (Fig.1C).

To test the achievable robustness and resolution of our workflow we used two benchmarks. First, the robustness of fractionation was assessed by the Pearson *R*^2^ between the two replicates of a given condition. Each condition showed an average correlation of *>*0.8 (Fig.1E), indicating high reproducibility in both phage infection and SEC fractionation, with most of the SEC-profile peaks overlapping within 1-2 fractions (*<*0.250 *µ*L). To test the resolution achievable with our chromatographic separations, we calculated the number of SEC peaks per proteins, which is a direct proxy for how many different complex assemblies a protein participates in. Approximately 45% of the identified proteins were detected in a single SEC-peak in each condition employed (Fig1F). While the presence of a single peak can represent detection of only a monomeric protein, we found the majority of these single-peak proteins are not at their predicted monomeric molecular weight (Sup. Fig.S1). This suggests that the protein complex assembly state of the PAO1 proteome was preserved during sample preparation and SEC fractionation.

### A high-quality interaction dataset for bacterial protein complexes

Next, we sought to investigate the recovery of known protein complexes by leveraging the partial conservation of core molecular assemblies between *P. aeruginosa* and other bacteria such as *E.coli*, for which protein complexes are more extensively annotated[21]. To visualize our data, we utilized the KZ resistant mutant dataset and projected it using t-SNE (Fig.2A). Smaller enzymes such as metabolic enzymes are usually co-expressed within the same operon[22] and have been reported to dimerize or multimerize. In line with this, we observed enzymes such as the pyruvate dehydrogenase complex (Fig.2B) and the oxoglutarate dehydrogenase complex (Fig.2C), which migrated at an estimated MW of *≈* 3.5 *∗* 10^6^ Da (expected MW *≈* 3.75 *∗* 10^6^ Da) and *≈* 2.4 *∗* 10^6^ Da, respectively. It is important to point that out that the molecular weight estimation for these large assemblies is subject to error due to these peaks being outside the external calibration curve. To achieve MW estimation we included in the calibration curve a pure SEC-separated 70S ribosome (Supplementary Fig. S2).

**Fig. 2.**
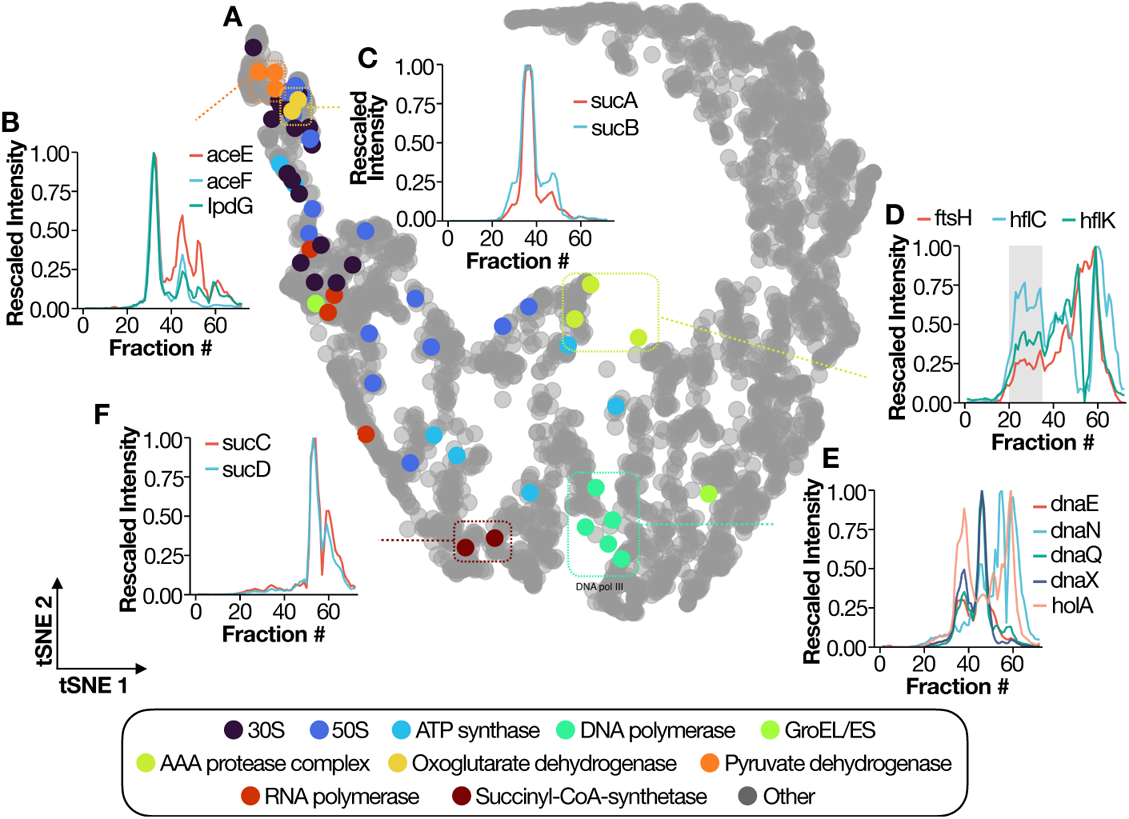
*Pseudomonas* protein complexes identified in the SEC-MS data. **A**. t-SNE plot for all the *P. aeruginosa* proteins detected in the KZ-res m1 strain. Each dot represents an individual protein, while the color code represents membership in reported protein complexes. Representative coelutions are showed for the pyruvate dehydrogenase complex (**B**), Oxoglutarate dehydrogenase complex (**C**), AAA protease complex (**D**), DNA polymerase III (**E**) and succinyl-coA synthetase (**F**). X axis shows the fraction number, while Y axis indicates the unit-rescaled intensity. Line color shows the various subunits.

Our sample preparation also preserved membrane-bound complexes. As an example, the AAA protease complex, formed by four hexamers of the AAA protease (ftsH) and 12 copies of each single-pass membrane proteins (HflK and HflC)[23], was recovered at high molecular weight in a broad peak as shown in Fig.2D. The large molecular weight range and sensitivity covered by our separation approach was also demonstrated in the recovery of more transient complexes such as the DNA polymerase III (dnaA, dnaE, and dnaQ) loaded with the *γ* complex (holA and dnaX) which plays a key role at the replication fork[24] (Fig.2E). Finally, heterodimeric complexes such as the succinyl-coA synthetase were also recovered as demonstrated by the coelution plot in (Fig.2F). Our manual inspections further confirm that prior knowledge can be easily incorporated into SEC-MS data analysis and allows for straight-forward identification of protein complexes.

### Comparison of host-targeted processes reveals conserved and divergent predation mechanisms

After having demonstrated the proteome depth achieved in our SEC-MS dataset and the recovery of known complexes, we turned our attention to how Jumbophages re-wire *P. aeruginosa* protein complexes by evaluating differences in SEC profiles upon phage infection. Variation in SEC profiles between conditions can arise from differential assembly state (i.e. a protein profile shifting to higher or lower molecular weight), different stoichiometry within a complex, or global alterations in protein abundance.

To quantify these different cases, we employed a previously described Bayesian analysis module from the PCprophet package[11] to derive marginal likelihoods (SEC differential score) of protein-level SEC changes between *ϕ*KZ and *ϕ*PA3 versus the receptorless infected samples (i.e. KZ-res m1). Comparing the SEC-profile differences between phage-infected PAO1 and KZ-res m1 revealed approximately 600 proteins showing SEC variation upon infection by either phage (Fig.3A). Notably, there is substantial consistency in which *P. aeruginosa* proteins are altered between both phages and the degree of change in their individual SEC profiles (Fig.3B, cor = 0.677), potentially pointing towards common pathways and complexes hijacked by *ϕ*PA3 and *ϕ*KZ for successful predation. Most of the changes at the assembly state level do not have a corresponding variation in protein abundance at the global proteome level, suggesting that SEC-MS offers an orthogonal view on effect of perturbations, such as infection, on the proteome (Fig.3C).

**Fig. 3.**
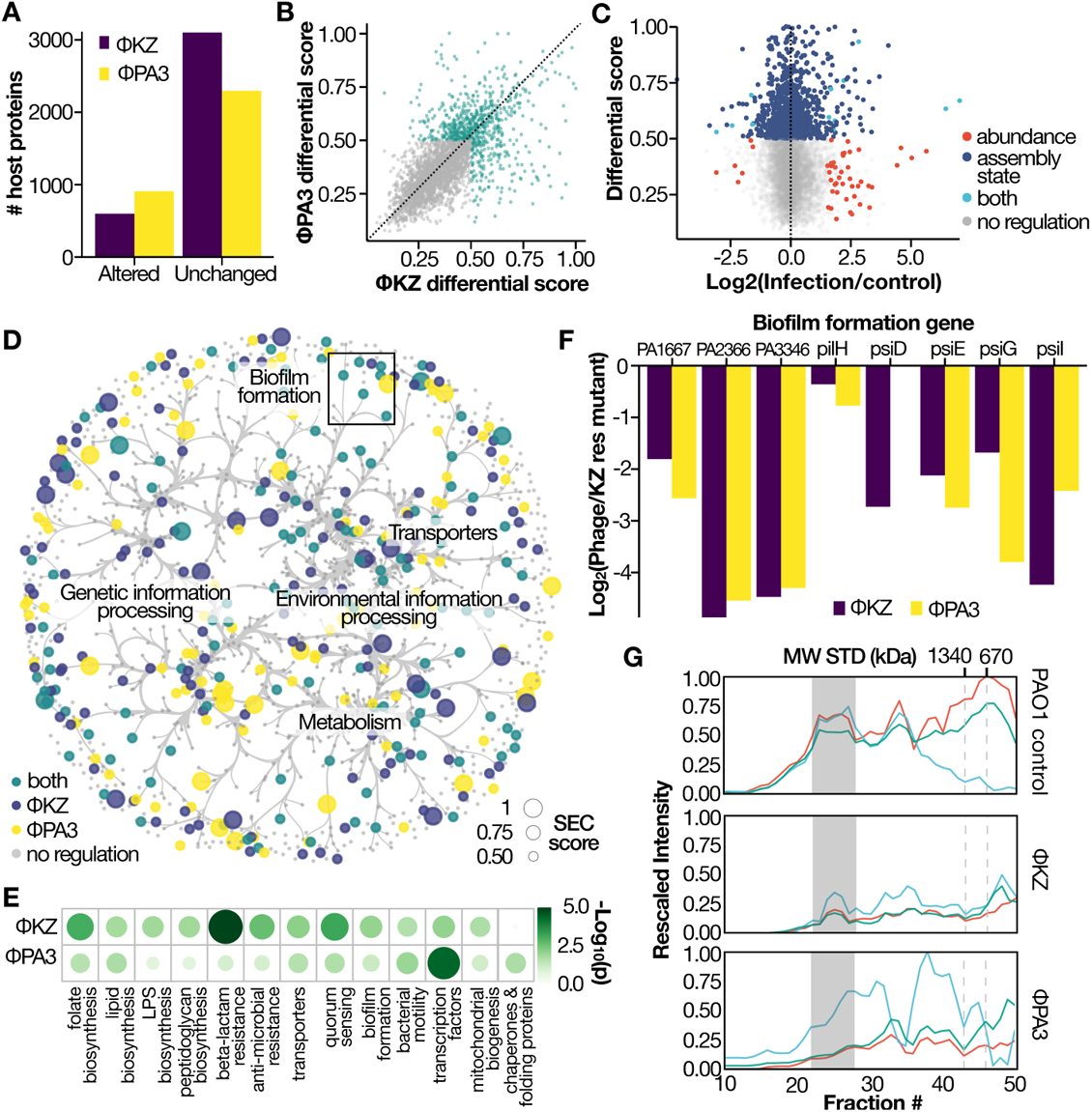
Differential analysis of SEC-MS data. **A**. Altered host proteins for each experiment. **B**. Scatterplot of differentially regulated proteins. Axis represents the differential SEC score while each dot represents a PAO1 protein. Color indicates significant regulation in either phage. **C**. 2D plot of differential SEC score (Y axis) vs LogFC from global proteome abundance (X axis). Color represents the different regulation level. Proteins highlighted in green are significantly regulated at the abundance level (Log2FC *≥* 2 and *q ≤* 1%) and assembly state level (SEC score *≥* 0.5). Both phages are shown. **D**. SEC derived molecular network for PAO1 proteins. Node color represents the regulation status, while node size shows the SEC score (i.e differential score). Edges are bundled using KDEEB. **E**. Enriched KEGG terms for altered proteins. Node size and color represents the significance on a −log10 scale. **F**. Barplot representing the log2FC biofilm formation genes upon phage infection as compared to control. KZ-res m1 experiments for the assembled MW range (i.e 2x monomeric weight for each protein). **G**. Coelution plot for the efflux pump MexA/B-oprM. Different experiments are represented by the various subpanels.

To identify conserved KZ-like jumbophage manipulation of the host interactome, we mapped the SEC-derived PAO1 interaction network (Fig.3D) with the correspondent protein-level differential data derived from the comparison between phage and uninfected samples. Although a large portion of the nodes do not have a functional annotation, we identified several classes where their components were significantly altered upon Jumbophage phage infection as depicted in Fig.3E. For example, the biofilm formation pathway (KEGG id:pae02025) was enriched in both phage infected samples (*q ≤* 0.01). Several prior studies have highlighted the role of phages in regulating formation of biofilms via modulation of polysaccharide production and perturbation cell envelope biology[25, 26]. We identified multiple proteins in this category having significantly decreased abundance in the high molecular weight region compared to their uninfected counterpart (Fig.3F), suggesting lower assembly state or complex reduction upon infection. Specifically, we observed *>*2 fold reduction in pslD, pslE and pslG in the high-molecular weight region. These proteins are members of a complex spanning from the inner membrane (pslG) to the outer membrane (pslD) which is required for the biosynthesis of exopolysaccharide[27]. The uncharacterized proteins PA3346, PA2366 and PA1667, display a broad coelution profile across the molecular weight dimension in the control sample, which is typically associated with membrane proteins[11]. Notably, these peaks are largely depleted (*>*3 fold reduction) in the infected condition. Outer membrane proteins were particularly affected by Jumbophage infection with porins and multi-drug efflux proteins (KEGG pae02010: ABC transporters) displaying significant reduction in interactions. For example, the MexAB-OprM complex, a key efflux pump[28], shows almost complete reduction of the fully assembled complex (Fig.3G). Importantly the MexAB-OprM complex was previously shown to be targeted by a Jumbophage closely related to *ϕ*KZ, called OMKO1[29], potentially representing a secondary receptor-binding site for *ϕ*KZ-like Jumbophages.

It is important to point out that changes we observed could either be beneficial for the phage to overcome its host, a host response to limit phage development, or simply be the result of pleiotropic regulators.

### Organization of *ϕ*KZ-like Jumbophage viral interactomes

The remodeling of host protein complexes can be the result of indirect rewiring of host cellular processes or direct interactions with phage proteins. Thus, we next investigated interactions directly involving phage proteins, including complexes containing both phage-host and phage-phage interactions.

Following SEC-MS and PPI prediction, we defined high-confidence interactions as those with a probability score of *≥* 0.75. In total, we identified 292 interactions between pairs of *ϕ*KZ viral proteins and 6550 host-pathogen interactions between *ϕ*KZ and PA01 proteins. *ϕ*PA3 showed a similar trend with 145 viral-viral and 3979 host-pathogen protein interactions (Fig.4A). Topological analysis of these networks revealed a scale-free architecture (Fig.4B), in line with previous reports that SEC-MS derived networks present the same architectural features as networks derived from literature curated studies and large PPI databases[11, 30, 31]. It has been observed with smaller phages, that genes within the same operon are often functionally related[32]. Accordingly, we evaluated the distribution of our predicted PPIs (by SEC-MS) in phage infected PAO1 cells as a function of the genomic separation of their corresponding genes (Fig.4C). Here we find a wide variation in the genomic distance between phage proteins that interact with other phage proteins (i.e. phage-phage interactions)(Fig.4C), with some genes being separated by distances as large as 139 kb. As example, the two RNA polymerases in *ϕ*KZ are both composed by proteins expressed in different operons with a max distance of 112.761 kb (PHIKZ080-PHIKZ180 in the vRNAp). Thereby, our resulting PPI distribution confirms the general lack of synteny within the genomes of *ϕ*KZ-like Jumbophages and shows the SEC-MS approach is a particularly advantageous technique to query phage encoded protein complexes, agnostic to the overall genome organization (i.e. a guilt-by-association approach at the protein level).

**Fig. 4.**
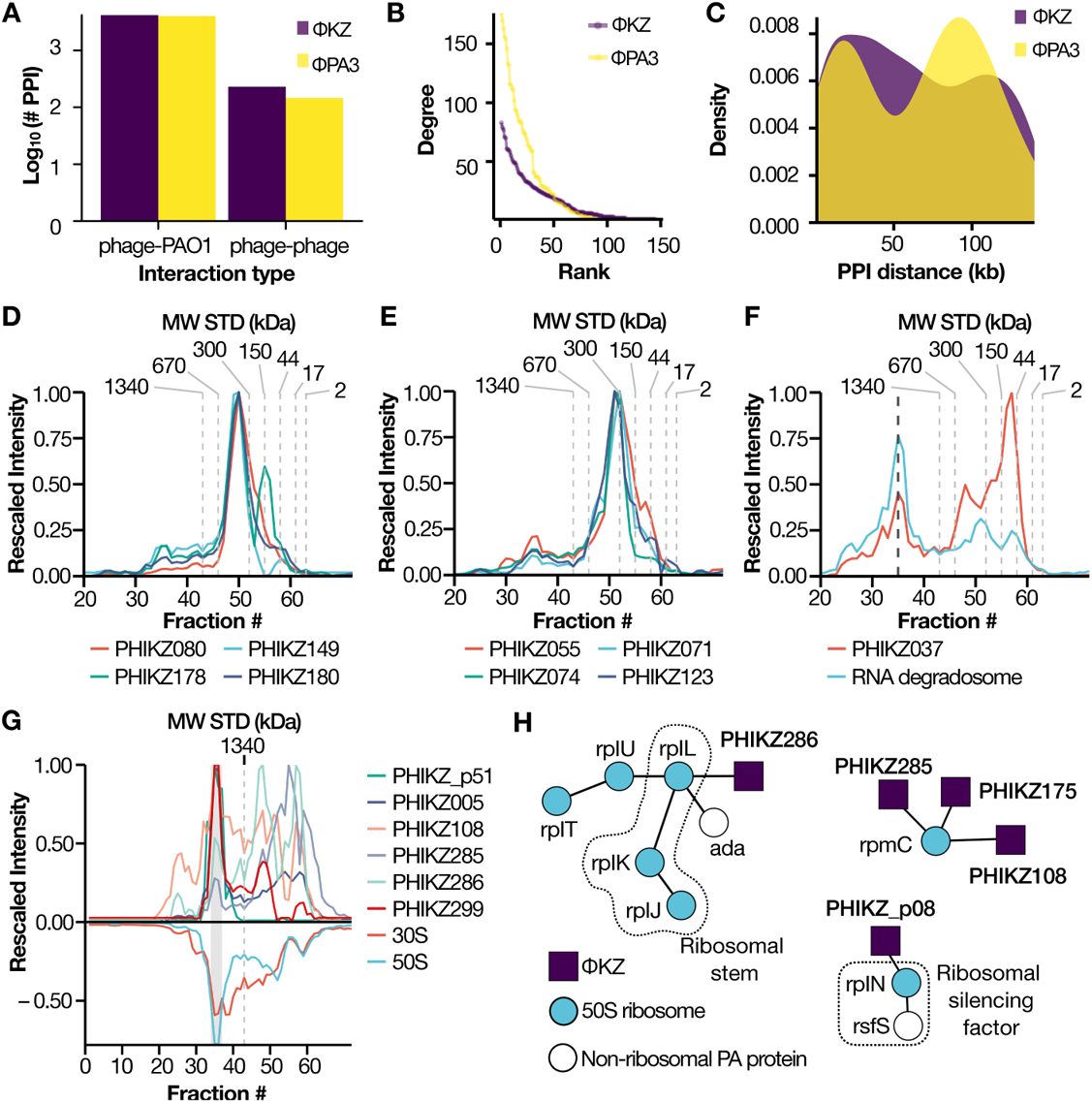
Comparative analysis of *ϕ*KZ and *ϕ*PA3 interaction networks. **A**. Number of PPIs identified for each phage (Y axis), separated by interaction type. **B**. Degree distribution (Y axis) versus the rank for a particular node (X axis). Line color represents the various phages. **C**. Density plot illustrating the genomic distance in Kb (X axis) for the phage-phage interactions. Color codes represent the two phages. **D, E**. Coelution plot for the *ϕ*KZ non-virion associated RNA-polymerase and the virion associated RNA-polymerase. **F**. Coelution plot for the RNA degradosome (cyan) and PHIKZ037 (red). Degradosome intensity represents the average across all six identified subunits (rhlB, deaD, pnp, rhlE, hfq, rne). **G**. Mirrorplot illustrating the coelution of *ϕ*KZ proteins (upper panel) with the 70S ribosome (lower panel). **H**. Interaction network for *ϕ*KZ proteins and ribosomal subunits from the SEC XL-MS experiment. Edges represent an identified crosslink. Dashed line represents a known ribosomal structural component, node color represents whether a protein is a phage protein (blue), ribosomal component (purple) or a non-ribosomal *P. aeruginosa* protein (white).

### Data-driven identification of *ϕ*KZ-like Jumbophage protein complexes

The identified interactions allowed us to recapitulate several known complexes in the Jumbophage proteome, despite a limited number being described at present. For example, we recovered the non-virion associated RNA-polymerase[33] migrating at its expected molecular weight (apparent MW 271 kDa, correct MW *≈* 265 kDa) (Fig.4D) as well as the virion associated RNA polymerase[34] (apparent MW 300 kDa, correct MW *≈* 297 kDa) as shown in Fig.4E. Previously described interactions between the phage and the host were also recovered by our approach, such as the interaction of PHIKZ037 with the RNA degradosome, which is involved in the accumulation of viral RNA[35] (Fig4F).

Building on the recovery of known phage protein complexes and the presence of several phage peak groups at high-molecular weight (Supplementary Fig.S3), we sought to probe our dataset for novel shell-associated proteins. Only two proteins so far have been identified as fundamental for shell formation and function: the major shell protein PhuN[36, 37] (gp54 in *ϕ*KZ; gp53 in *ϕ*PA3, gp105 in 201-*ϕ*2-1) which is the main building block of the shell complex[14, 16, 38], and the bipolar tubulin spindle protein phuZ which serves to stabilize the shell in the center of the bacterial cell[39]. Because the capsid docks on the shell prior to tail attachment and lysis[17], we are unable to differentiate phage proteins contained within the capsid from those that are ejected and associated with the shell by apparent increases in molecular mass alone. To distinguish between these two cases, we performed two orthogonal control experiments using a cesium purified virion sample and a shell-enriched sample (Supplementary Fig. S4A), which we used to filter the SEC-MS interactors to only proteins enriched in the shell sample and absent in the virion (Supplementary Fig. S4B-C). The six remaining proteins (PHIKZ p22, PHIKZ036, PHIKZ111, PHIKZ p64, PHIKZ232 and PHIKZ261) were then tested for association with the shell via fluorescence microscopy using PAO1 expressing mNeonGreen tagged constructs. Overexpression of most constructs resulted in diffuse localization outside of the phage shell (Supplementary Fig. S4D), with gp36-mNG and p64-mNG displaying puncta that were sometimes peripheral to the phage shell, but timelapses revealed them to be mobile throughout the cell (see Supplementary Data). Of note, these results do not conclusively exclude these proteins as potential shell components. Further confirmation of these results would be needed to address potential technical issues, such as disruption of protein localization by tagging or over-expression, shell association at an earlier or later stage of infection, or difficulty in detecting transient interactions. In addition, large (*>*MDa) and intact shell fragments have been shown to be mostly insoluble[37], likely resulting in loss of many shell fragments prior to SEC and tightly bound shell-associated proteins as a result.

We then turned our attention to interactions between phage and host complexes. Interestingly, several *ϕ*KZ proteins (PHIKZ005, PHIKZ108, PHIKZ285, PHIKZ286, PHIKZ299, and PHIKZ p51) were predicted by our deep learning tool to be in complex with the fully assembled *P. aeruginosa* 70S ribosome (Fig.4G). To validate these *ϕ*KZ proteins as ribosomal interactors, we performed cross-linking mass spectrometry (XL-MS)[40] on a pooled sample from the SEC fractions corresponding to the 70S ribosome (Supplementary Fig.2). We identified 975 crosslinks in total (202 hetero-links and 871 homo-links), covering several previously reported bacterial protein complexes (Supplementary Fig.S5). The XL-MS data recovered 24 *P. aeruginosa* ribosomal proteins (separated in 30S and 50S) of which 3 showed physical interaction with 5 *ϕ*KZ proteins. Amongst these phage proteins, we recovered PHIKZ285, PHIKZ286, and PHIKZ108, which were predicted from the SEC-MS data to be in complex with the 70S ribosome. Moreover, we identified PHIKZ p08 and PHIKZ175 as additional ribosomal interactors (Fig.4H). PHIKZ286 bound the L1 ribosomal stalk (rplL, rplK, and rplJ) which has an important role in tRNA translocation[41] and is the contact site for several translation factors[42]. PHIKZ p08 interacted with rplN bound to its ribosome silencing factor rsfS which slows down or represses translation[43]. Finally, PHIKZ285, PHIKZ175, and PHIKZ108 were bound to rpmC which is an accessory protein positioned near the exit site and required for triggering nascent polypeptide folding[44]. These findings demonstrate the power of SEC-MS to detect HPIs involved in critical aspects of host biology, however further mechanistic characterization is needed to determine if such phage proteins manipulate ribosomal, or instead represent active translation of the phage proteins.

### Identification of novel proteins by SEC-MS

The multiplexed nature of DIA allows un-biased sampling of the full precursor space[45], so we queried our data for the presence of peptides from novel phage proteins using a custom protein FASTA built with EMBOSS. We detected 4 previously undescribed proteins for *ϕ*KZ (2 forward and 2 reverse ORFs) and 11 for *ϕ*PA3 (8 forward and 3 reverse) (Fig.5A and B). The authenticity of these novel proteins is supported by the detection of two or more proteotypic peptides for nearly all proteins (Fig.5C) and reproducible detection of the same peptides in 15 or more consecutive fractions across independent experiments (Fig.5D). All novel proteins showed reproducible quantitation between biological duplicate experiments (n=72 per replicate), with an average protein-level correlation of 0.75 for *ϕ*PA3 proteins and 0.82 for *ϕ*KZ proteins (Fig.5E). Most of these proteins migrated at a higher molecular weight than their predicted molecular weight, suggesting they may be associated with high-order assemblies (Fig.5F). Some of the novel ORFs are further supported by a great degree of sequence overlap with homologs. The most staggering example is the *ϕ*PA3 reverse sense ORF 56450-58417 which shows *>*70% sequence similarity with previously reported proteins from various *Pseudomonas* spp. phages (*ϕ*KZ, Psa21, Phabio, 201*ϕ*2-1, and PA1C)(Supplementary Fig. S6A). Interestingly, all proteins showing *≥* 50% identity to 56450-58417 are previously reported or proposed phage RNA polymerase components (RNAP), such as PHIKZ074 (non-virion associated RNAp, UniprotID Q8SD88)[34, 46, 47]. To date, there is no experimental evidence of a nvRNAP in *ϕ*PA3. To derive other putative members of this complex, we extracted the predicted interactors of ORF 56450-58417 (Fig 5G) and performed BLASTp analysis to identify proteins showing homology to other Jumbophage RNA polymerase components. From this analysis, we selected 3 interactors (AVT69 gp055, AVT69 gp063, and the novel ORF 53811-55010) showing *>*50% conservation with multiple Jumbophage proteins annotated as RNAP components (Supplementary Fig.S6B-D). Specifically, we identified homologs of both the *β^′^* polymerase subunit (AVT69 gp055 and ORF 56450-58417), as well as homologs of the *β* subunit (ORF 53811-55010). The *ϕ*PA3 protein AVT69 gp063 displays 57% sequence similarity to PHIKZ068, an essential nvRNAp component which lacks structural similarity to know components of previously reported RNA polymerases[46]. Utilizing the position of the SEC peak, we estimated the nvRNAp MW in *ϕ*PA3 to be *≈* 321 KDa (Fig.5G). Assuming the lack of homodimers in the structure, the predicted MW for these four proteins was *≈* 253 KDa, suggesting a putative missing subunit. Of note, we did not identify 53811-55010 interactors corresponding to PHIKZ123, another *β* subunit component, which could explain this observation. To explore the possibility of these proteins (AVT69 gp055, AVT69 gp063, ORF 53811-55010, and ORF 53811-55010) folding into an RNA polymerase-like assembly we performed structural prediction of this peak group using AlphaFold2 multimer[48]. We aligned the best scoring model (ipTM + pTM = 0.82, Fig.S7) to the reported structures for the *ϕ*KZ nvRNAP (PDB 7OGP and 7OGR)[47] as depicted in Fig.5H. We reached a template modeling (TM) score of 0.503 using US-Align[49] and an average RMSD of 1.016 Å using MatchMaker[50] between our proposed *ϕ*PA3 vRNAp and the *ϕ*KZ RNAp (70GR), indicating a shared tertiary structure similarity between these two assemblies. As we obtained low distances for the *β* and *β^′^* subunits, we set to investigate the misaligned region at the Cterm of the polymerase clamp (AVT69 gp063 in *ϕ*PA3 and PHIKZ068). Despite showing high sequence homology (68%), these two proteins share a large intrinsically disordered region (IDR) in the middle of the sequence (275293 aa for PHIKZ068 and 277301 aa AVT69 gp063) as shown in Supplementary Fig.S8). The IDR likely enables flexibility in the central region, resulting in varied orientation for the folded C-term in AVT69 gp063 following AlphaFold predictions.

**Fig. 5.**
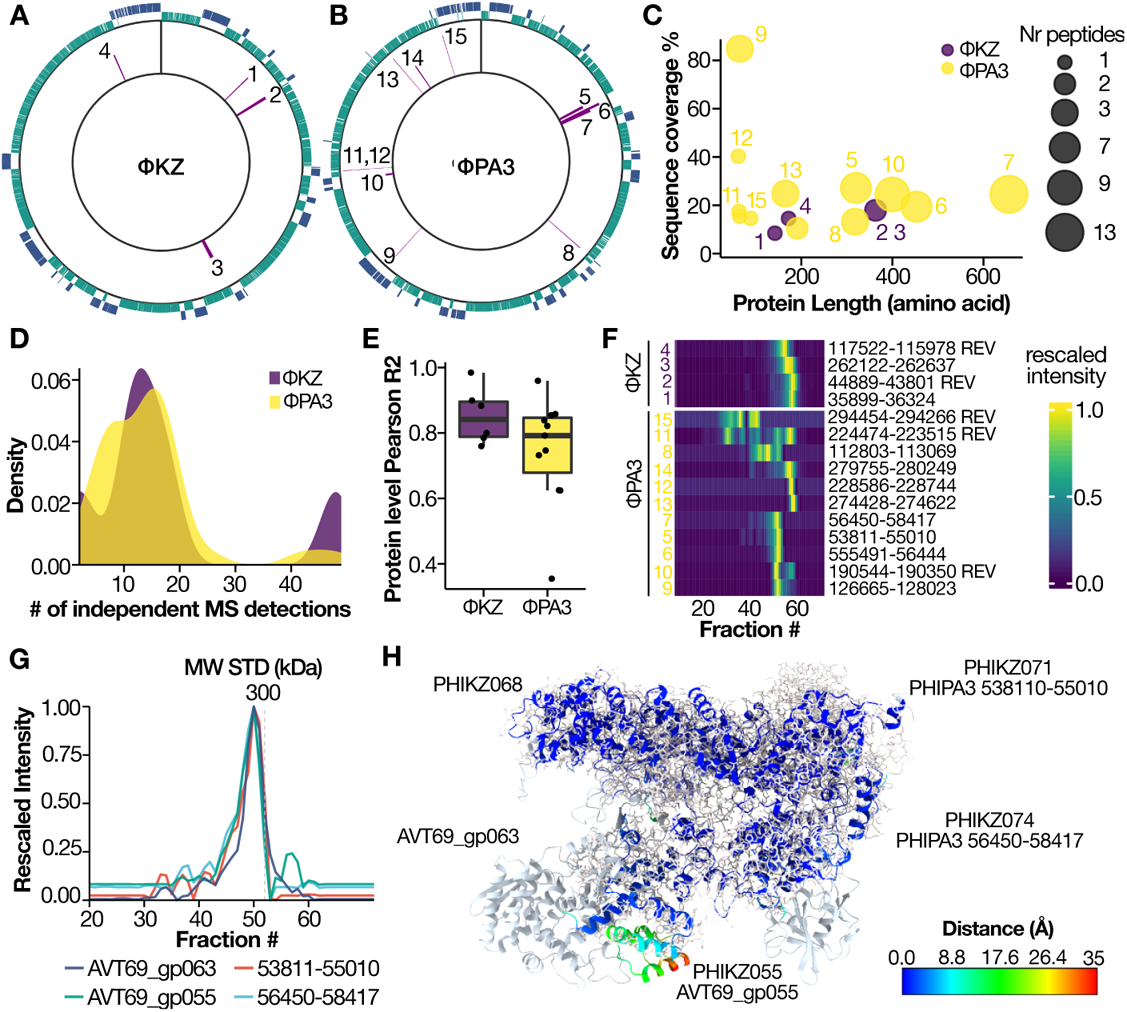
Identification of novel phage proteins. **A-B**. CDS plot for *ϕ*KZ and *ϕ*PA3. Forward CDS are colored in green while reverse CDS are represented in purple. Identified novel proteins are highlighted in the histogram (inner circle). **C**. Scatterplot of protein length vs percentage of sequence coverage in the SEC-MS experiments. Dot size represents the number of proteotypic peptides identified. **D**. Distribution of the identifications (defined as number of independent MS detections using a 1% peptide spectrum matching FDR) for the novel *ϕ*KZ and *ϕ*PA3 proteins. **E**. Boxplot showing the Pearson correlation between the two replicates (n=72). Each novel protein is represented as a dot. The box boundaries show the interquantile range (IQR) and its whiskers 1.5*×*IQR. **F**. Heatmap representing the elution profile for all the novel ORFs. X axis represents the fraction number while the cell color shows the unit-rescaled intensity. **G**. Coelution profile for predicted nvRNAp in *ϕ*PA3. **H**. Superimposition of reported structure for the *ϕ*KZ nvRNAp (stick, grey) and predicted structure for the *ϕ*PA3 nvRNAp (ribbons). Chains are colored by their distance to the *ϕ*KZ nvRNAp structure after superimposition.

### Discovery and validation of novel injected phage proteins

Although it is well known that *ϕ*KZ phages guard their genome from nucleolytic host-immune systems by building a proteinaceous shell[14], this structure is only visible after 20 minutes of infection. Little is known about how the phage genome is protected or packaged prior to shell assembly. To identify phage proteins proximal to the genome, with possible protective functions, we first determined a detailed virion proteome to allow distinction of virion proteins (injected) from newly synthesized proteins. We performed cesium chloride-gradient purification of *ϕ*KZ coupled with deep peptide fractionation and long chromatographic acquisition (see Supplementary Methods for details). The 245 *ϕ*KZ proteins identified in this dataset encompassed *≥* 90% of previously reported head proteins (Supplementary Fig S.9A). To account for low-level contamination caused by cesium chloride-fractionation, we compared our enriched virion sample with the previously reported virion proteins to derive an ROC curve, which we used to select an intensity threshold maximizing recall of known virion proteins and minimizing false positive rate (Fig. 6A). This filtering identified 81 significantly enriched proteins in total. This included 58/61 (95%) of those previously reported and added 23 proteins to the virion composition which are strongly enriched over their corresponding protein abundance in a non-enriched samples (Fig. 6B). This drastic increase in protein number is dependent on the increased sensitivity sequencing speed of the MS utilized for acquisition as well as extensive offline sample fractionation prior to MS acquisition (see Supplementary Fig S. 9B).

**Fig. 6.**
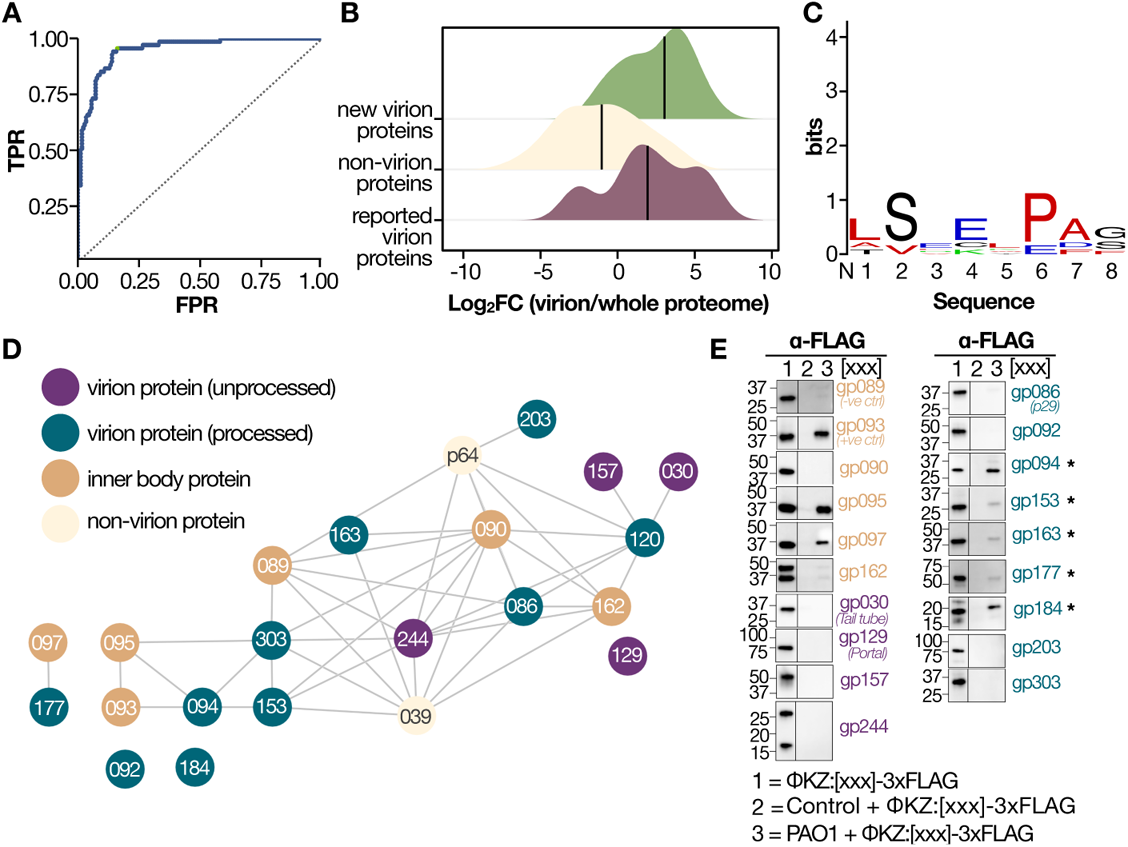
Data-driven analysis of injected inner body proteins. **A**. ROC curve for virion MS (AUC *≈* 0.94) using as ground truth the prior reported virion proteins. Green highlight selected threshold for maximum sensitivity at the lowest FPR. **B**. Density plot representing the enrichment of virion proteins over a whole proteome infection experiment expressed as log2 fold change (X axis). Different colors represent whether a protein was previously reported as virion (bordeaux), novel from our virion dataset (green) or non-virion (cream). **C**. Sequence logo for the proteins in the IB interaction network. X axis shows the position from N to C term while Y axis represents conservation in bits. **D**. SEC-MS derived interaction network for the reported IB proteins (gp93/95/97). Color code represents the query protein (aquamarine), abundant virion protein (dark purple) defined as top 20% most abundant proteins, non abundant virion proteins (grey) and proteins not identified in the virion MS experiment (green). **E**. Injection of phage proteins evaluated by western blot of 3x FLAG phage tagged proteins. gp089 (-ve ctrl) serves as a negative control, while gp093 (+ve ctrl) serves as a positive control. Validated novel injected proteins are designated with an asterisk.

As prior work reported extensive gp175-driven proteolysis of the head and inner body (IB) proteins[51], we searched our purified virion data for evidence of unexpected termini (i.e. semi-tryptic peptides) with the goal of confirming prior reported cleavages and potentially identify novel ones. We recovered 63 semi-tryptic peptides, of which 15 could be mapped to prior data[51] (*≈* 40% overlap). Within our semi-triptic peptides, we identified 20 cleavages corresponding to the reported IB proteins (gp93/95/97) and 12 mapping to 9 unreported proteins. Of note, 8 cleavages could be mapped to gp94, gp177, and gp303 (see Supplementary data and Supplementary Fig. S9C). To identify consensus sequences within our set of IB interactors we performed motif enrichment analysis using STREME[52]. We found that the LSxE consensus motif was enriched (BH-adjusted *p* = 6*e^−^*^5^), corroborating the previously reported one S/A/G-X-E motif, while providing additional specificity in the P2 position (Fig6C).

Starting with only the virion proteome, we next queried the *ϕ*KZ interaction network to identify putative injected proteins (Fig. 6D). Building on this data, we selected the interactors of the previously reported proteins (gp94, gp153, gp162, gp163, and gp177) for additional validation using our previously reported assay for evaluating injection[18]. In this assay, PAO1 cells expressing the protein of interest with a FLAG tag are infected with wild-type *ϕ*KZ, resulting in phage particles with labelled proteins. These phage particles are then used to infected gentamyicin treated PAO1 cells (i.e cells where translation is inhibited) allowing to evaluate injection using a western blot. By further lowering the interaction thresholds to positive predicted interactions (i.e PPI probability *≥* 0.5 instead of 0.75 utilized to select high-confidence interactors), we further identified gp184 as an IB interactor and validated it as an injected protein. These experiments, confirmed injection of the previously reported IB proteins (gp93, gp95, gp97) and further validated the injection of most their interactors (gp94, gp153, gp163, gp177, gp184) as showcased in Fig.6E. We note that among the 3xFLAG-tagged virion proteins tested in this study, the injection profile of inner body protein PHIKZ090 is inconsistent with a previous report[18]. In this prior study, it was reported that PHIKZ090 tagged at the C-terminus with mNeonGreen is injected into PAO1 cells upon infection by PHIKZ (as detected by fluorescence microscopy). We do not detect injection of PHIKZ090-3xFLAG via western blotting. The reasons for this discrepancy are unclear and may result from weaker expression of this construct (as compared to the controls PHIKZ093-3xFLAG and PHIKZ089-3xFLAG) and lower levels of packaging in the virion. As a consequence the amount of PHIKZ090-3xFLAG injected into PAO1 cells is reduced, likely to levels that are below the detection limit of the western blot assay. We acknowledge that poor expression of certain constructs is a limitation of this assay. Here, by using an highly sensitive MS of the virion combined with SEC-MS, we identify and validate the injection of eight proteins (three previously reported) that are highly abundant, found in the virion, and interact with the previously reported IB proteins. Overall, these proteins give us a starting point to unravel the interactome of the ejected phage genome and identify proteins that protect the genome from host nucleases.

## Discussion

Understanding the dynamics driving host and pathogen interactions and their dynamics upon infection is a crucial component to deepening our knowledge on the mechanisms regulating infection progression and outcome. To date, most proteomics studies of infectious diseases focused on the analysis of a few pathogen proteins by tag/antibody-based purification or the measurement of protein abundance variation in infected samples. Yet it is widely known that the pathogen proteome works as an ensemble through protein-protein interactions to hijack the host cell which in turn regulates both expression and interaction between host proteins. Hence, a system-wide view on the intrinsic modularity of the pathogen proteome and how it quantitatively regulates host complexes is key to understanding pathogenic mechanisms at the molecular level.

In this study we demonstrate the first application of SEC-MS to systematically investigate pathogen proteome organization and host interactome plasticity upon Jumbophages infection of *P. aeruginosa*. *ϕ*KZ-like phages (specifically *ϕ*KZ and *ϕ*PA3) are potent killers of *P. aeruginosa* (with a broad host range), making them timely alternatives to antibiotics with many *ϕ*KZ-like phages already in clinical trials to treat bacterial infections. By obtaining an atlas of these phages interactomes, we can begin to construct a mechanistic understanding of the *ϕ*KZ-like Jumbophage infections, ranging from viral composition to protein injection, transcription, and phage nucleus assembly and growth.

Our *ϕ*KZ-like phage interactomes recapitulated prior evidence for the subdivision of Jumbophage proteomes into distinct assemblies such as virion and non-virion associated RNA polymerases, as well as the interaction with key host complexes such as the RNA degradosome. We expanded our knowledge on the phage interactions with essential host processes such as translation where we identified phage proteins interacting with the ribosomal stem and ribosomal silencing factors. Moreover, while the lack of immediate genome organization hinders the prediction of functions for phage proteins, the deep coverage and unbiased nature of SEC-MS data offers a straightforward approach to identify novel complexes and propose putative functions. As an example, by using SEC-derived interactors of a de-novo predicted *ϕ*PA3 protein (ORF 56450-58417), we identified an heterotetrameric assembly which is predicted to have strong structural homology to the reported nvRNAP in *ϕ*KZ. This suggests that the unbiased nature of SEC-MS data allows for not only the discovery of an uncharacterized protein, but also enables to probe its putative function through the detection of new protein-protein interactions. Identifying such complexes will enable further investigation using structural and biochemical approaches. In addition to the identification of interactions, these maps offer the opportunity to further quantify host interactome remodelling and disentangle variation in expression from assembly state. By comparing the *P. aeruginosa* interactome between infected and uninfected, we observed a large degree of changes during infection, with perturbation of similar complexes between the two Jumbophages suggesting conserved mechanisms of phage predation. While here we a first draft of the KZ-like Jumbophage interactome, it is important to acknowledge the trade-off between specificity and throughput in interaction identification in SEC-MS, which we mitigated by utilizing only high-confidence interactions for analysis. Advances in deep learning models for prediction of interactions from co-fractionation mass spectrometry data and integration of orthogonal features (beside the coelution itself) such as predicted structure or function is expected to improve prediction accuracy and reduce the false discovery rate for uncharacterized proteomes. Overall, the characterization of host-pathogen molecular networks remains challenging, but we provided the first interactome-wide study of infection progression using two models *ϕ*KZ-like phages in *P. aeruginosa*.

Wider application of SEC-MS is expected to significantly accelerate the characterization of pathogenic mechanisms by providing proteome-wide insights into the physical association between host and pathogen complexes, thus enabling identification of novel druggable targets, host vulnerabilities, or guidance in the development of novel biologicals.

## Data availability

The supporting MS data is available via ProteomeXchange with the identifier PXDXXXX. The PhageMap database is freely accessible at https://phagemap.ucsf.edu/ and the underlying interactions have submitted to the BioGRID database and STRING database. Novel *ϕ*PA3 and *ϕ*KZ proteins have been submitted to UniProt. All the code to reproduce the plots as well as the intermediate data and Alphafold2 predicted structures are available on GitHub at https://github.com/anfoss/Phage data.

## Acknowledgments

This work was supported by an NIH grant 1R01AI167412 (JBD and DLS) and 1R01AI171041 (JBD and DAA). We thank Prof. James Wells at UCSF for the usage of the HPLC used to perform the size-exclusion experiments. Molecular graphics were performed with UCSF ChimeraX, developed by the Resource for Biocomputing, Visualization, and Informatics at the University of California, San Francisco, with support from National Institutes of Health R01-GM129325 and the Office of Cyber Infrastructure and Computational Biology, National Institute of Allergy and Infectious Diseases. Figure 1A-B was prepared with Biorender.

## Author contributions

AF: Performed proteomics sample preparation and analysis all MS data, developed PhageMAP and wrote the manuscript. DLS, JBD, DA, NK: Conceptualization, supervision, writing, and funding acquisition. AP: Novel ORF prediction DM. CK. BG: Phage infection experiments, virion enrichment, microscopy and WB for injected proteins EM, MM: Performed shell enrichment YP: Critical input in revising the manuscript. All co-authors contributed in reviewing and editing the manuscript.

## Supplementary figures

**Supplementary Fig. S1.**
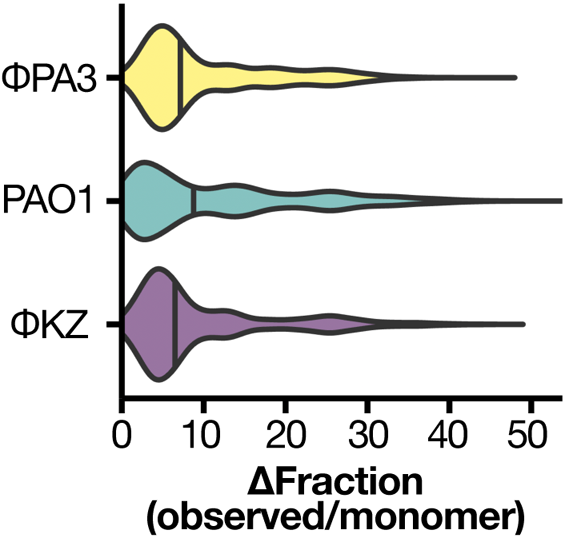
Violin plot showing the distance between the observed protein SEC peak and their predicted molecular weight expressed as fraction number for the single-peak proteins. Black line represents the mean.

**Supplementary Fig. S2.**
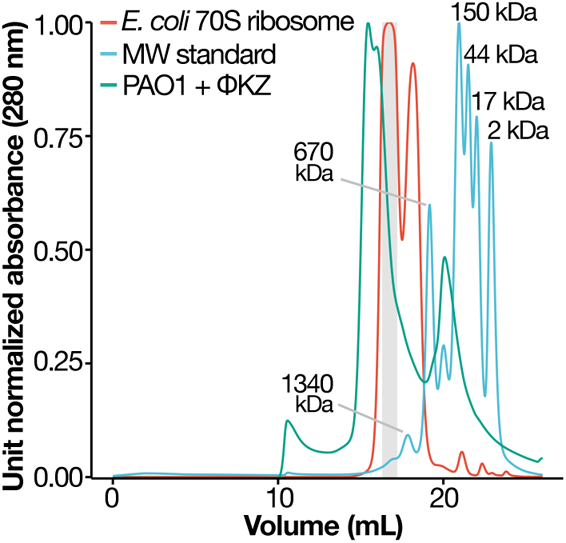
Ribosome absorption profile and MW from SEC. Chromatographic traces collected at 280 nm for the purified 70S ribosome (red), the protein mixture used as molecular weight standards (cyan) and the *ϕ*KZ infected sample (green). The grey area shows the fractions used for the ribosomal crosslinking experiment.

**Supplementary Fig. S3.**
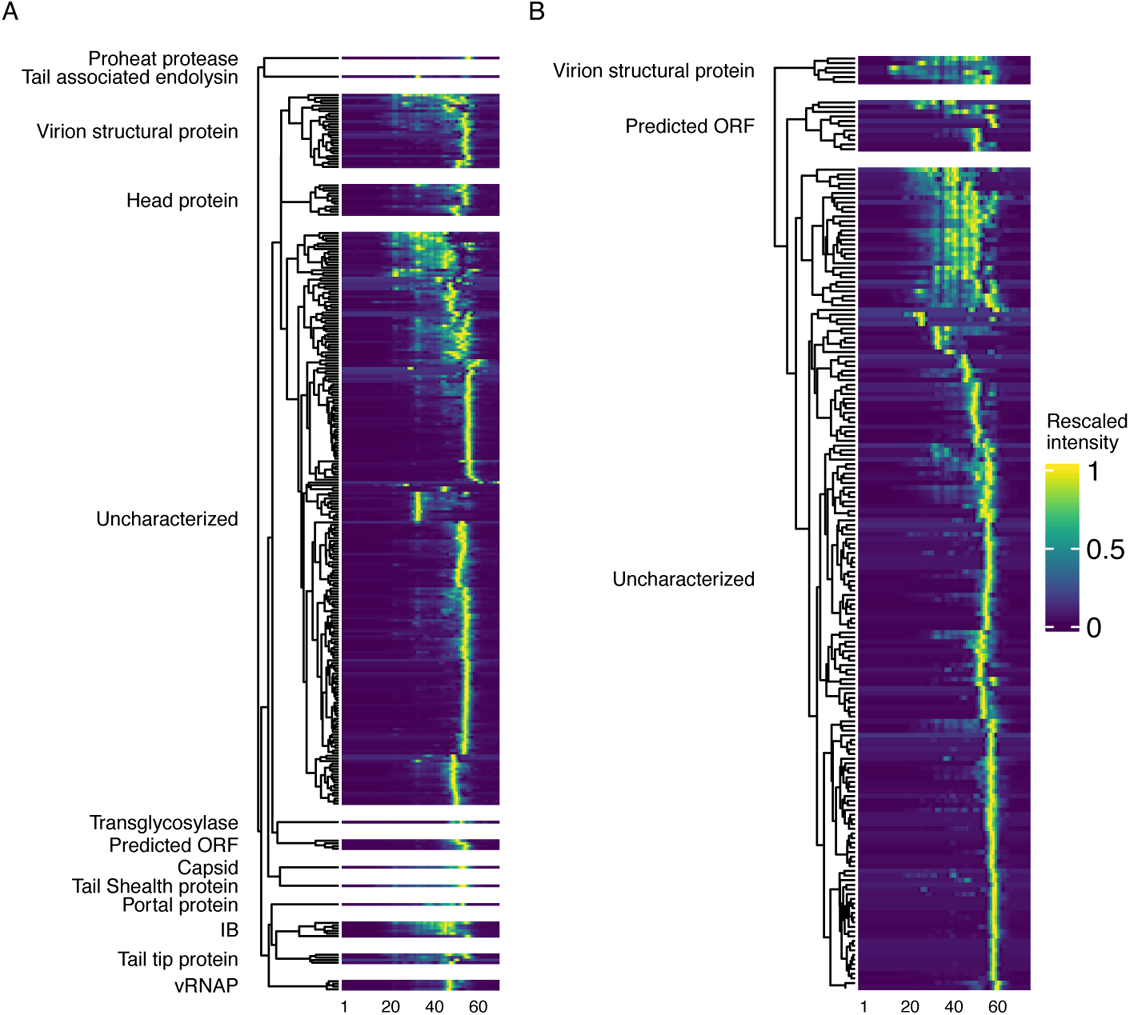
Distribution of the KZ-like phages proteomes into discrete assemblies. **A-B**. Coelution heatmap for all the phage proteins identified in *ϕ*KZ (**A**) and *ϕ*PA3 (**B**). The dendrogram branches are labelled based on manual literature curation for the corresponding proteins in the peak group. Color represents the unit-rescaled intensity. X axis represent the fraction number.

**Supplementary Fig. S4.**
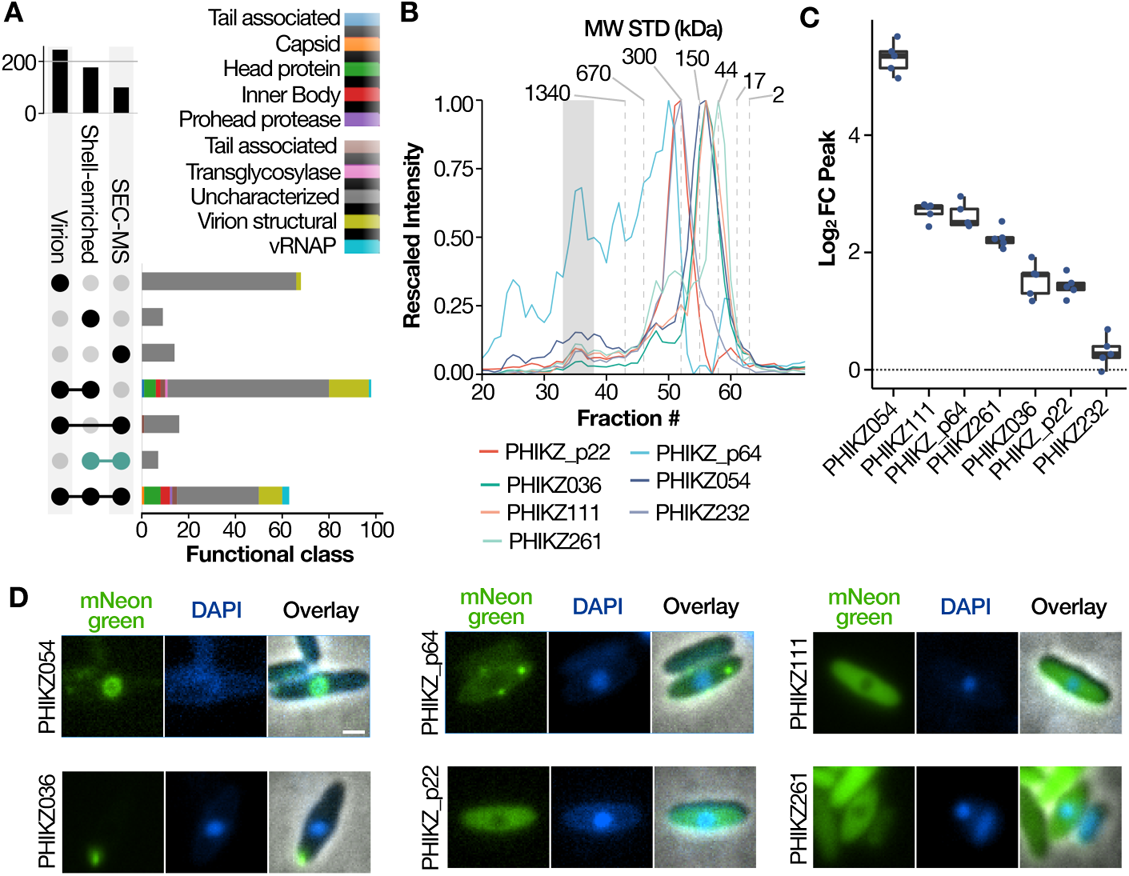
Predicted *ϕ*KZ shell proteome. **A**. Upset plot showing the overlap between the *ϕ*KZ virion enrichment, shell enrichment and SEC-MS experiment. Green barbell shows the retained putative shell proteome **B**. Coelution profile for main shell protein (gp54) and its predicted interaction partners. **C**. Enrichment (Log2FC) over average protein intensity for the proposed shell proteins D. Fluorescent microscopy of proposed shell associated proteins.

**Supplementary Fig. S5.**
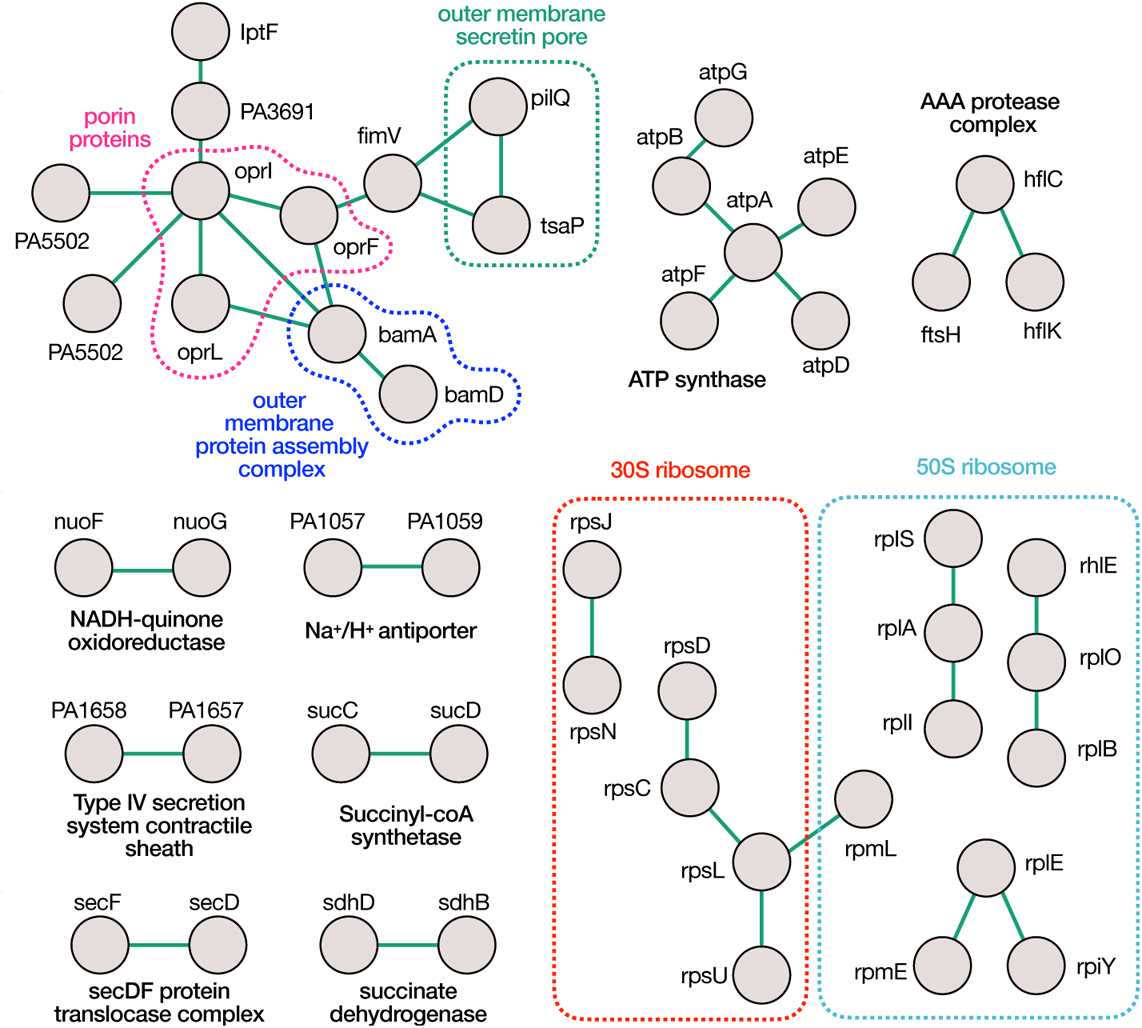
Know protein complexes recovered by SEC-XL MS. Interaction network derived from the heterolinks detected at 5% CSM-FDR for reported complexes in *Pseudomonas*. Text label shows the complex for the interacting proteins.

**Supplementary Fig. S6.**
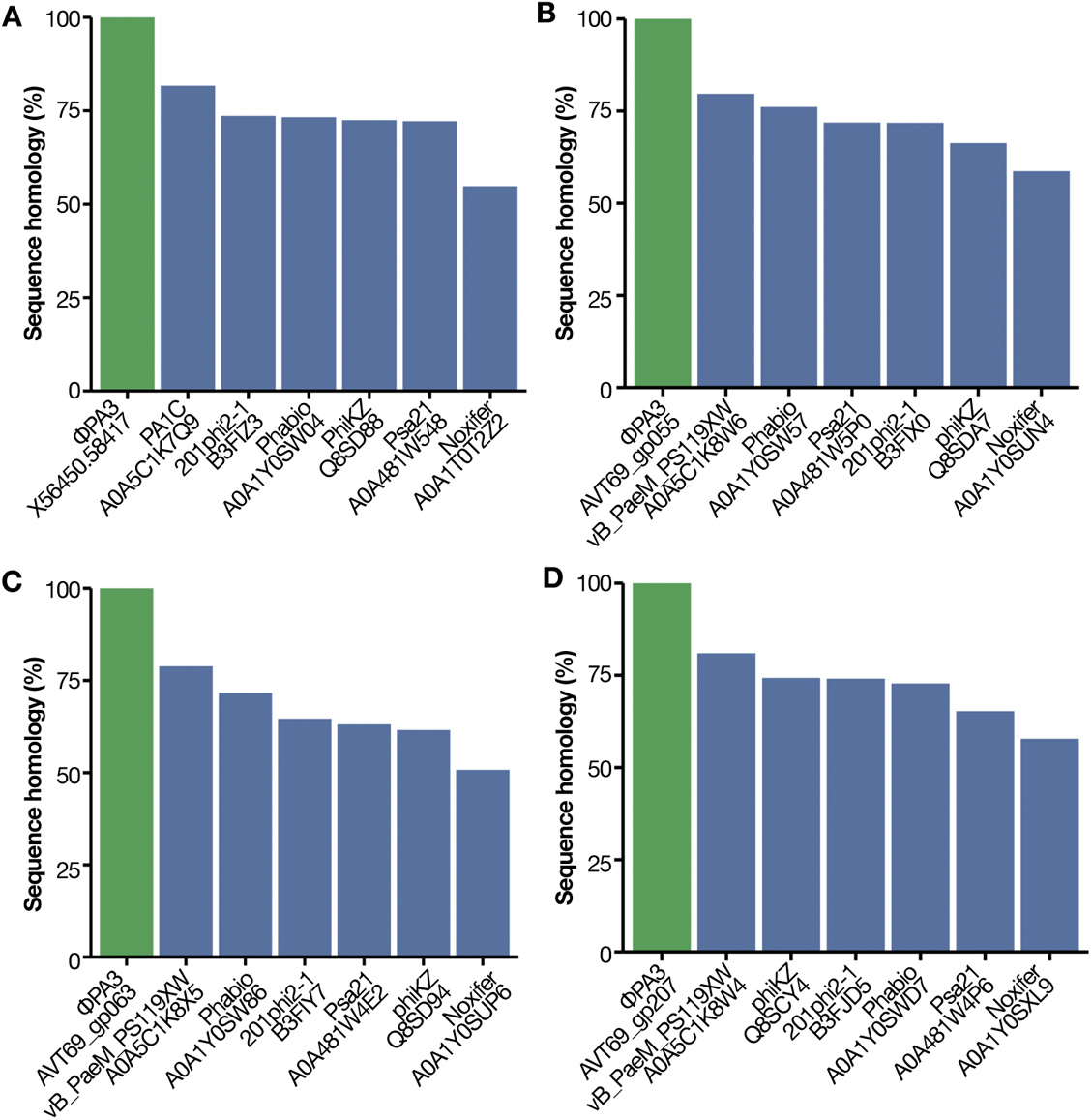
Alignment of *ϕ*PA3 56450-58417 interactors with other Jumbophage proteins. **A-D**. Barplot showcasing the sequence homology between ORF 56450-58417 (**A**), gp55 (**B**), gp63 (**C**) and gp207 (**D**) to other *Pseudomonas* phages protein. *ϕ*PA3 proteins are highlighted in green. Y axis shows the percentage of sequence homology.

**Supplementary Fig. S7.**
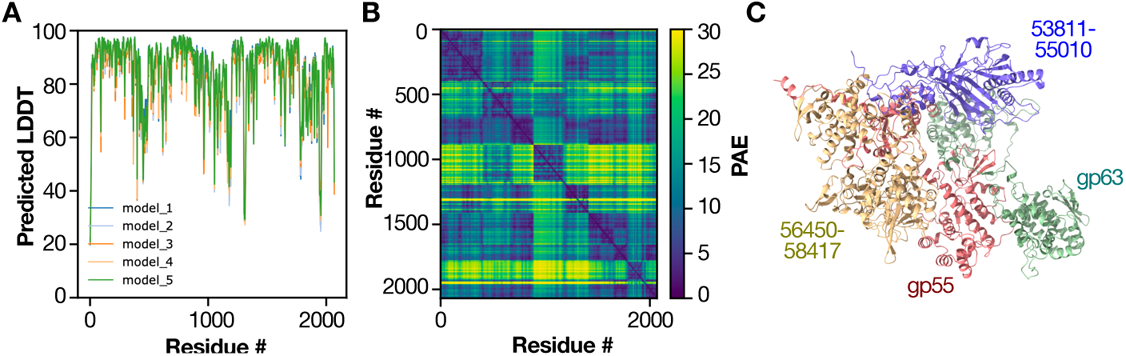
Prediction of *ϕ*PA3 non virion associated RNA structure. **A**. Per-residue local confidence (pLDDT) versus sequence (X axis). Different line colors represents the different AF2 model **B**. Predicted alignment error (PAE) heatmap **C**. Structure of best scoring model (iPTM + TM = 0.826)

**Supplementary Fig. S8.**
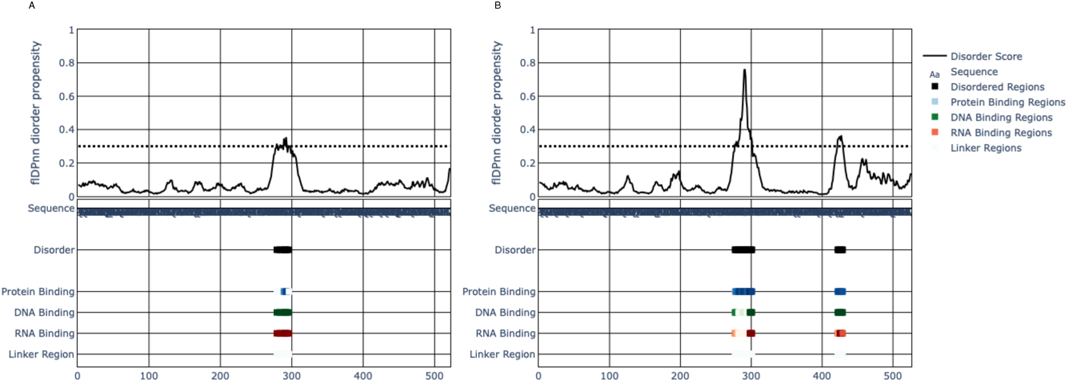
Intrinsically disordered region in *ϕ*KZ gp68 and *ϕ*PA3 gp63. Prediction of disordered regions using flDPnn[53]. X axis represent sequence, while different rows shows different local predicted properties between *ϕ*KZ gp68 (**A**) and *ϕ*PA3 gp63 (**B**)

**Supplementary Fig. S9.**
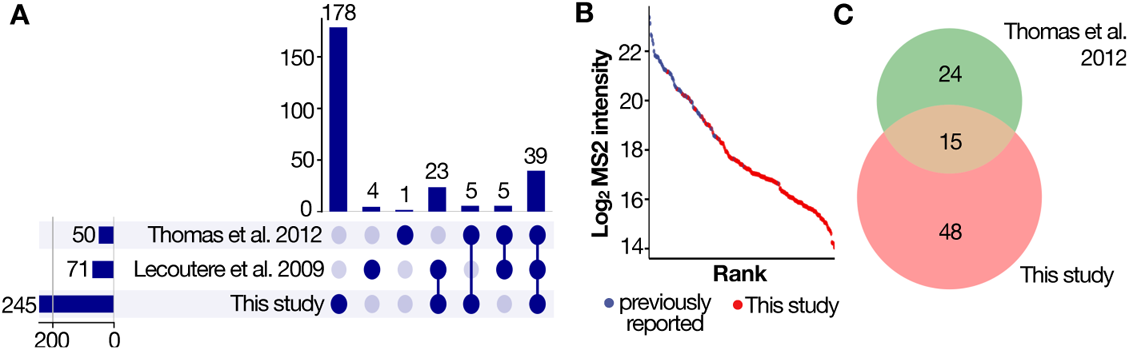
Benchmark of purified virion MS versus other reported studies of *ϕ*KZ head proteins. **A**. Upset plot showing the overlap in protein IDs between this study, Thomas et al. (2012)[51] and Lecoutere et al. (2009) [54] **B**. Distribution of intensities for virion proteins identified in this study. Blue proteins were previously identified while red proteins are novel virion proteins from this study. **C**. Venn diagramm of semi-tryptic peptides detected in this study versus Thomas et al[51]

## Supplementary tables

**Table 1.**
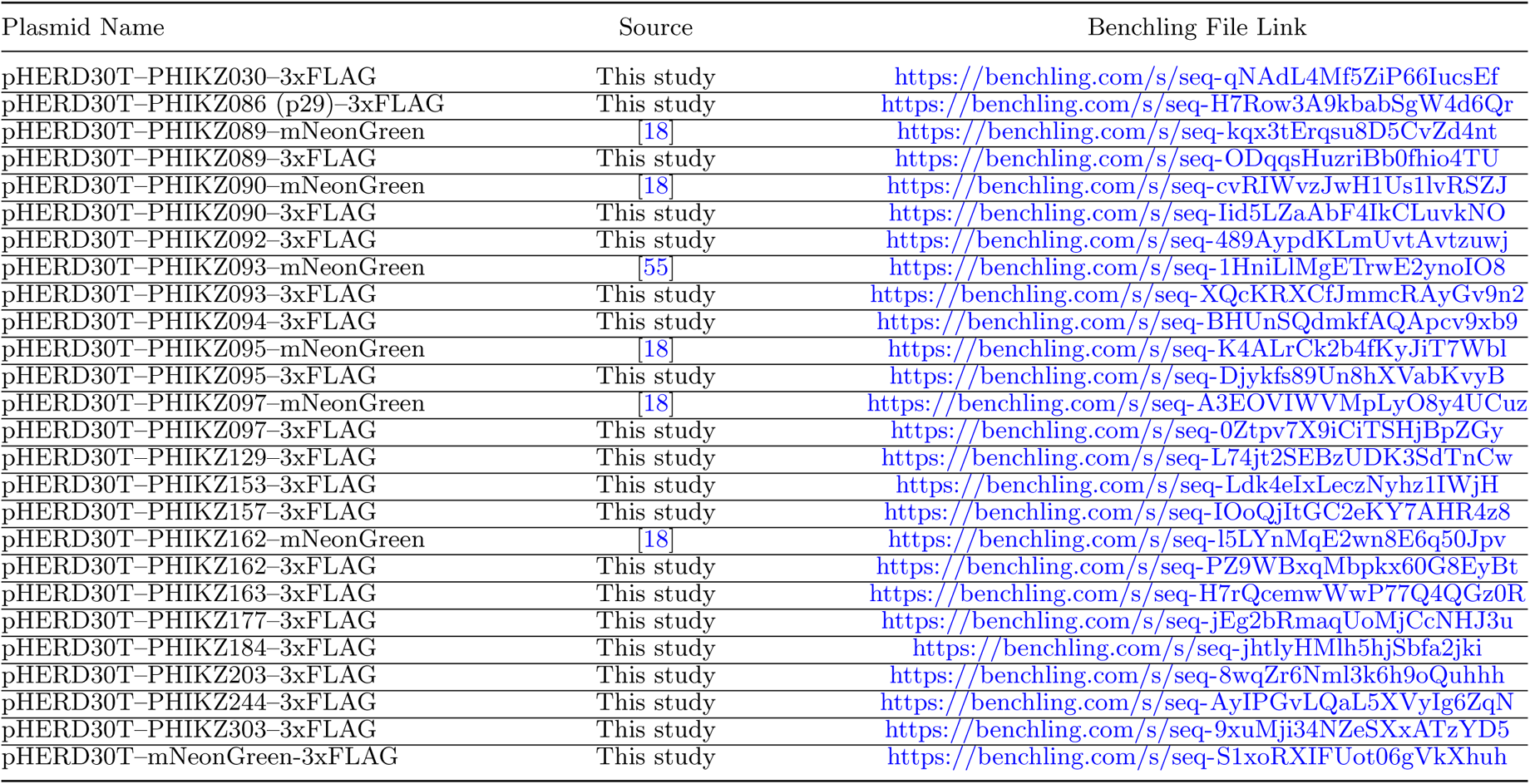
Plasmid sequences utilized in this study

**Table 2.**
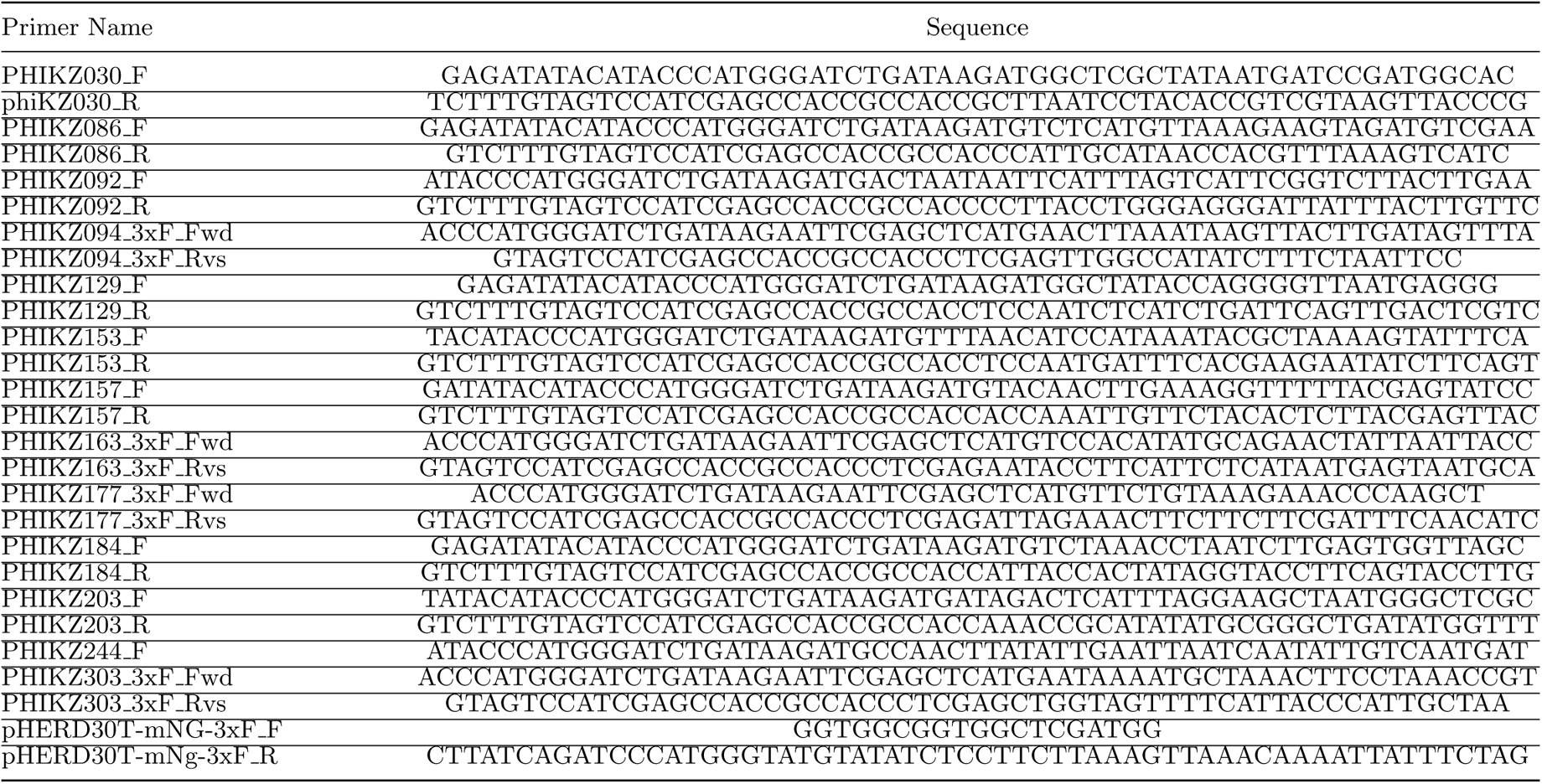
Primers utilized in this study

## Material and methods

### Cloning

C-terminal 3xFLAG fusions of *ϕ*KZ IB proteins: pHERD30T plasmids encoding C-terminal 3xFLAG fusions of *ϕ*KZ IB proteins (PHIKZ089, PHIKZ090, PHIKZ093, PHIKZ095, PHIKZ097, PHIKZ162) were cloned as follows: the plasmids pHERD30T-IB-mNeonGreen (PHIKZ089, PHIKZ090, PHIKZ093, PHIKZ097, PHIKZ162, sequence: Supplementary Table 1) were digested with restriction enzymes XhoI and KpnI (upstream and downstream of the mNeonGreen sequence) to generate a pHERD30T-IB backbone (IB: PHIKZ089, PHIKZ090, PHIKZ093, PHIKZ097, PHIKZ162). An insert sequence corresponding to XhoI-GGGGS-3xFLAG-KpnI was digested with XhoI and KpnI to generate an insert fragment that was ligated individually with each corresponding backbone using T4 DNA ligase to generate the plasmids. The pHERD30T plasmid encoding a C-terminal 3xFLAG fusion of PHIKZ095 was generated as follows – The plasmid pHERD30T-PHIKZ090-3xFLAG was digested with restriction enzymes SacI and XhoI to generate a pHERD30T-3xFLAG backbone. The plasmid pHERD30T-PHIKZ095mNeonGreen (sequence: Supplementary Table 1) was digested with SacI and XhoI, and the smaller fragment corresponding to SacI-PHIKZ095-XhoI was extracted by DNA Gel extraction to obtain an insert fragment. The insert fragment was ligated with the pHERD30T-3xFLAG backbone using T4 DNA ligase to create the plasmid pHERD30T-PHIKZ095-3xFLAG. All plasmid sequences were checked for correct assembly upstream and downstream of the insert sequence with Sanger Sequencing (Quintara Biosciences) using sequencing primers QB0068 (5*′*–ATGCCATAGCATTTTTATCC–3*′*) and QB0049 (5*′*–CCCAGTCACGACGTTGTAAAACG–3*′*). Cterminal 3xFLAG fusions of *ϕ*KZ virion proteins: pHERD30T plasmids encoding C-terminal 3xFLAG fusions of *ϕ*KZ proteins (PHIKZ030, PHIKZ p29, PHIKZ092, PHIKZ094, PHIKZ129, PHIKZ153, PHIKZ157, PHIKZ163, PHIKZ177, PHIKZ184, PHIKZ203, PHIKZ244, PHIKZ303) were generated as follows: the plasmid pHERD30T–mNeonGreen –3xFLAG (Supplementary Table 1) was linearized via PCR using primers pHERD30T-mNG-3xF F, pHERD30T-mNg-3xF R (Supplementary Table 2) to generate a pHERD30T –3xFLAG backbone (removing the mNeonGreen sequence). The genes encoding *ϕ*KZ proteins were amplified from purified *ϕ*KZ particles (removing the stop codon) via PCR using the corresponding primers (Supplementary Table 2) to generate insert fragments. These insert fragments were individually assembled with the pHERD30T –3xFLAG backbone to generate the plasmids. All plasmid sequences were checked for correct assembly upstream and down-stream of the insert sequence with Sanger Sequencing (Quintara Biosciences) using sequencing primers QB0068 (5*′*–ATGCCATA GCATTTTTATCC–3*′*) and QB0046 (5*′*–TGTAAAACGACGGCCAGT–3*′*). Benchling files containing sequences of all constructs, attached primers and sequencing files are reported in Supplementary Table 1.

### Bacterial culture

*Pseudomonas* aeruginosa strains PAO1 were grown overnight in 3 mL LB at 37°C with aeration at 175 rpm. Cells were diluted 1:100 from a saturated overnight culture into 100 mL LB with 10mM MgSO_4_ and grown for *≈* 2.5 hours at 37°C with aeration at 175 rpm. At OD600nm = 0.5-0.6 (*≈* 3*e*^8^ CFU/mL), the cell cultures were infected with bacteriophage (*ϕ*KZ or *ϕ*PA3; MOI *≈* 1) on ice for 10 minutes (to allow complete adsorption of virions onto cells) and then incubated at 30°C for 50 minutes (total time of infection 60 minutes). Thereafter, the cell cultures were transferred to pre-chilled 50 mL falcon tubes, centrifuged at 6000x*g*, 0°C for 5 minutes. The supernatant was discarded and cell pellets were washed twice with 5 mL ice-cold LB and combined. After the final wash, the bacterial pellets were resuspended in 5 mL ice-cold LB. The concentrated cell culture was flash frozen in liquid nitrogen and subsequently mechanically lysed using a SPEX-freezer mill.

### Shell isolation via density centrifugation

The shell isolation was performed as we previously reported[37]. Briefly, we infected *P. aeruginosa* PA01 with *ϕ*PA3 or *ϕ*KZ for 60 minutes. The bacteria was mechanically lysed via Dounce homogeneization in NP40 Lysis Buffer (50 mM Bis-Tris, 150 mM NaCl, 0.5% NP40, 5% glycerol, 5 mM DTT, 20 ng/*µ*l Lysozyme, 1 mM EDTA, 1mM EGTA – pH 6.5). The lysate was clarified at 16000x*g* for 5 min and the insoluble fraction resuspended in wash buffer (20 mM Bis-Tris, 150 mM NaCl, 1 mM DTT, 1 mM EDTA, 2 mM MgCl2 – pH 6.5). This was subject to further 500x*g* (5 min) and 15000x*g* (10 min) centrifugation with the insoluble fraction isolated and resuspended in wash buffer each time. The insoluble fraction of the last 15,000x*g* spin was retained as the final product. The shell enriched sample was acetone precipitated using 8 volumes of ice-cold acetone and incubated overnight. Following incubation, the protein pellet was washed thrice with ice-cold acetone and dried under vacuum.

### Cesium gradient purification of phage virions

Bacteriophages (*ϕ*KZ or *ϕ*PA3) were propagated in LB at 37°C with PAO1 as a host. Liquid growth curve experiments were used to ascertain the MOI of bacteriophage stock needed to ensure complete lysis of the bacteria following a substantial growth as ascertained by OD600 measurement. Growth curve experiments were carried out in a Synergy H1 micro-plate reader (BioTek, with Gen5 software). Cells were diluted 1:100 from a saturated overnight culture with 10 mM MgSO_4_. Diluted culture (140 *µ*l) was added together with 10 *µ*l of 10X serial dilutions of bacteriophage stocks to wells in a 96-well plate. This plate was cultured with maximum double orbital rotation at 37°C for 24 h with OD600 nm measurements every 5 minutes. Thereafter, the bacteriophage stock was added at the appropriate MOI to a 1:100 back-dilution of a saturated PAO1 overnight culture in 100 mL LB with 10mM MgSO_4_ and the bacterial culture incubated for 24 hours (37°C with aeration, 175 rpm). 5 mL of chloroform was added to the cultures in a fume-hood and the cultures were incubated to with chloroform for 15 minutes (37°C, 175 rpm) to ensure maximum lysis of bacterial cells. The cell cultures were transferred to 50 mL falcon tubes and centrifuged at 6000 x*g* for 15 min to pellet bacterial debris. The supernatant (containing bacteriophages in high titer) was carefully transferred to a fresh set of 50mL falcon tubes and centrifuged and 6000x*g* for 15 min to pellet any residual bacterial debris. The supernatant was transferred to fresh 50 mL falcon tubes with 2 mL chloroform. To obtain high purity virion particles, a previously described protocol was followed[56]. The virions from the bacterial cell lysate were concentrated by slow stirring overnight at 4°C in 1 M NaCl and 10% PEG (final concentration) and then pelleted (11’300x*g*, 4°C, 30 min). Pellets were resuspended in 20 ml of SM buffer (50 mM Tris-HCl (pH 7.5), 100 mM NaCl, 8 mM MgSO_4_, 0.002% gelatin) containing Complete Protease Inhibitor (Roche). The phage suspension (5.8 mL/tube) were layered onto CsCl step gradients composed of the following concentrations of CsCl: 1.59 g/ml (0.75 ml), 1.52 g/ml (0.75 ml), 1.41 g/ml (1.2 ml), 1.30 g/ml (1.5 ml) and 1.21 g/ml (1.8 ml). The buffer used throughout the gradient was 10 mM Tris-HCl (pH 7.5) and 1 mM MgCl2. Tubes were spun at 31,000 rpm for 3h at 10°C in an SW41 rotor (Beckman Coulter ultracentrifuge) and the resulting phage band had a buoyant density of 1.36 g/ml. This fraction was collected and dialyzed against three changes of 50 mM Tris-HCl and 10 mM MgCl_2_ at 4°C. This ultra-purified phage stock was diluted in SM buffer and its titer assessed using plaque assays. Finally, the phage virion stock was acetone precipitated using 8 volumes of ice-cold acetone.

### Bacterial infection and SEC sample preparation

Cryomilled samples were resuspended in *≈* 4 ml of SEC running buffer (50 mM ammonium bicarbonate and 150 mM NaCl pH 7.4) supplemented with protease inhibitors (Roche) and ultracentrifuged at 60’000 *g* for x minutes at 4°C. The supernatant was concentrated to 100 *µ*L using a 100 KDa molecular weight cutoff filters to simultaneously enrich for high-molecular weight assemblies and deplete monomeric proteins. The concentrated sample was centrifuged once more at 10’000 g at 4°C to remove particles.

### Size-exclusion chromatography

Approx 1000 *µ*g per sample (*≈* 80 *−* 90*µ*L as estimated by Bradford’s assay) were separated on a Agilent Infinity 1260 HPLC operating at 0.5 mL/minute in SEC running buffer with a Phenomenenex SRT-C1000 column connected and cooled at 4 °C. 72 fractions of 125 ul were collected after 3.75 ml until 13 ml and the column was then washed with 2 column volumes (18 mL) of SEC buffer. The MW was estimated using a protein mixture (Phenomenex AL0-3042), while a *E.Coli* 70s ribosome (NEB, cat nr P0763S) was used to estimate which fractions to use for ribosome XL-MS.

### SEC-MS proteomics sample preparation

The SEC samples were prepared as we previously reported[57] using a 96 well filter-aided sample preparation (FASP). The FASP-filters were conditioned by washing twice with 100 *µ*L of ddH_2_0. SEC buffer was removed by centrifugation (1800 *g* 1 h) and proteins were resuspended in 50 *µ*L of TUA buffer (TCEP 5 mM, Urea 8M, 20 mM ammonium bicarbonate) and incubated on a thermos shaker (37°C, 400 rpm) for 30 minutes. Cysteine residues were then alkylated by addition of 20 *µ*L CAA buffer (Chloroacetamide 35 mM, 20 mM ammonium bicarbonate) for 1 h at 25°C in the dark. TCEP and CAA were removed by centrifugation (1800 *g*, 30 min) and filters were washed 3 times with 100 *µ*L of 20 mM ammonium bicarbonate. Proteins were digested in 50 *µ*L of 20 mM ammonium bicarbonate with 1 *µ*g of tryspin per fraction. A 96 well receiver plate (Nucleon, Thermo-Fisher) was used to collect the peptides by centrifugation for 30 minutes at 1800*g*. The filter plates were washed once with 100 *µ*L of ddH_2_O and centrifuged to dryness (1800*g*, 60 minutes). The peptides from the receiver plate were transferred to protein LoBind tubes (Eppendorf) and the corresponding well was washed with 50 *µ*L of 50% acetonitrile (ACN) in ddH_2_0 to increase the recovery of hydrophobic peptides. The combined resulting peptides per each fraction were vacuum dried and stored at −80 C until MS-acquisition. For each phage, 5 *µ*L from each fraction were pooled together to generate a phage-specific library. Each sample specific library was prepared on a C18 spin column (Nest). Following activation of the column with 1 column volume (CV) 100% ACN and wash with 2 CV of 0.1% formic acid the peptides were bound to the column and eluted using a step-wise gradient of ACN from 5 to 25 (5% increases) in 0.1% triethylamine to account for the increased hydrophobicity of the XL peptides compared to not modified ones. A final fraction at 80% ACN was added to recover hydrophobic peptides.

### Proteomics sample preparation for virion enriched protein pellets

Dried proteins were resuspended in 100 *µ*L of 8M urea, 100 mM ammonium bicarbonate (ABC) pH 8.1. TCEP (Thermo Fisher) was added to 5 mM final concentration and the samples were incubated at room temperature for 30 minutes. Reduced cysteines were alkylated with 10mM chloroacetamide (CAA) for 30 minutes in the dark. Following alkylation, the urea was diluted to 1 M with 100 mM ABC and the proteins were digested with 2 *µ*g of trypsin per sample for 14 hrs at 37°C in a thermo-shaker (600 rpm). Digestion was stopped by acidification using 10% formic acid (FA) and the samples were desalted using a C18 spin column (Nest group). Briefly, columns were activated using 1 column volume (CV) of ACN and then equilibrated with 2 CV of 0.1% FA. Peptides were loaded twice and then washed with 3 CV of 0.1% FA. Elution was done using 0.5 CV of 50% ACN 0.1% FA and repeated twice. Samples were dried under vacuum and stored at −80°C until acquisition.

### Crosslinking MS sample preparation

*ϕ*KZ infection and SEC-separation were performed as described above. Following separation, the SEC-fractions corresponding to the 70S ribosome peak (F33-F38) were pooled. The was crosslinked for 1 hr at RT using 5 mM DSSO from a freshly prepared 30 mM stock in water-free DMF. The reaction was quenched by addition of ABC to 50 mM for 30 minutes at RT and the proteins were precipitated using 8 volumes of ice-cold acetone. Following overnight incubation, pellets were washed 5 times with 8x volumes of ice-cold acetone and briefly dried under vacuum. The pools were reconstituted in 8M urea, 100 mM ABC and 5 mM TCEP and incubated for 30 minutes at RT. CAA was added to 10 mM final concentration and the samples were incubated in the dark for 1 hr. Urea was diluted to 1 M by addition of 100 mM ABC and the proteins were digested overnight with 2 ug of trypsin in a thermo shaker at 30 °C. Samples were acidified with 10% TFA and high-ph tip fractionation was performed as we previously described[57]. Briefly, following activation, equilibration and washing of the C18 resin, the elution was done using a step-wise gradient of ACN from 10 to 40 (5% increases) in 0.1% triethylamine to account for the increased hydrophobicity of the XL peptides compared to not modified ones. Resulting fractions were dried under vacuum.

### SEC-MS and spectral library acquisition

Samples were resuspended in buffer A (0.1% FA) and approximately 200 ng were analyzed by DIA-PASEF on a Bruker TimsTOFpro interfaced with a Ultimate3000 UHPLC. For the SEC-MS experiment, the peptides were separated on a PepSep column (15 cm, 150 um IID) using a 38-minute gradient at 0.6 *µ*l/min. Following loading, the peptides were eluted in 20 minutes with a 5% to 30% B (0.1% FA in ACN) in 20 minutes. The column was then washed for 5 minutes at 90% and high flow (1 *µ*l/min) and re-equilibrated at 5% ACN for the next run. The peptides were sprayed through a 20 mm ZDV emitter kept at 1700 V and 200 °C. The mass spectrometer was operated in positive mode using DIA-PASEF acquisition[58]. Briefly, 4 PASEF scans (0.85 1*/K*0 to 1.30 1*/K*0) were acquired and divided each precursor range into 24 windows of 32 Da (500.7502 – 966.67502 *m/z*) overlapping 1 Da. Each of the fractionated samples (phage-specific libraries) was acquired in DDA-PASEF using a similar gradient composition except for the elution which was performed in 90 minutes leading to a 120 minute gradient. For DDA-PASEF the ion mobility window and precursor range were matched to the DIA boundaries to allow for seamless library building and search.

### XL-MS data acquisition

The XL-MS samples were acquired on a Bruker TimsTOFpro interfaced with a Ultimate3000 UHPLC. The peptides were separated using a 118 minutes linear gradient. Following loading, the percentage of B (80% ACN in 0.1% FA) was increased from 2% to 8% in 5 minutes and then to 43% in 90 minutes. Residual peptides were eluted at 50%B for 10 minutes and then the column was washed at 88% B for the remaining 13 minutes. The peptides were separated on a PepSep column (15 cm, 150 mm iid, 1.9 *µ*m beads size). The mass spectrometer was operated in positive mode and data-dependent acquisition with the same source parameters as the SEC fractionated samples. To enrich for crosslinked peptides a custom IM polygon was employed[59] and charge inclusion was enabled (3 + *to*8+ precursors). Precursors having nominal intensity above 20’000 were selected for fragmentation using an inverted collision energy of 23 *eV* at 0.73 1*/k*0 and 95 *eV* at 1.6 1*/k*0.

### SEC-MS data analysis

The DDA files were searched within the Fragpipe framework using MSfragger[60] and the ‘DIA-speclib-quant’ workflow using the *Pseudomonas aeruginosa* pan proteome FASTA (5564 entries, proteome ID UP000002438, downloaded on the 05/22). For each phage, the correspondent FASTA nucleotide file was downloaded from GenBank (NC 004629.1 for *ϕ*KZ and NC 028999.1 for *ϕ*PA3) and EMBOSS was used for novel ORFs prediction (see ‘Prediction of novel ORFs’ section for details). The GenBank files were translated to protein level using BioPython and supplemented to the *Pseudomonas* FASTA. Carbamylation of cysteines was set as fixed modification while oxidation of methionine, N-term acetylation (peptide level) and pyroglu formation were set as variable modifications. EasyPQP (https://github.com/grosenberger/easypqp) was used to generate a spectral library. Following phage-specific library generation, PAO1 precursors from all libraries were transferred to ensure the presence of the same PAO1 proteins with the same peptides across all DIA experiments using lowess for RT realignment. The DIA-PASEF data was searched with DIA-NN[61] v.1.7.1 using a library-centric approach. Identified spectrum with MS1 precursors within 10 ppm and MS2 precursor within 15 ppm were selected and a second library was generated (double-pass mode). Quantification was set to robust (high-accuracy) and cross-run normalization was disabled.

### XL-MS data analysis

XL-MS timsTOF files were converted to mgf using MSconvert. MS1 peak picking was enabled and the spectrum were denoised (top30 peaks in 100 m/z bins). Ion mobility scans were combined. Following the conversion, the peak files were searched in XiSearch[62] using a fraction-specific FASTA containing only the protein ids identified by SEC-MS in the corresponding MW range. MS1 and MS2 tolerances were fixed to 10 and 15 ppm with 10 ppm of peptide tolerance. DSSO was selected as crosslinker (158.0037648 Da) and the correspondent oxidized and amidated crosslinker were added as modifications. Link-FDR was fixed at 5% (boosted) and the resulting file were imported into XiView (https://xiview.org) for manual inspection of crosslinked spectrums.

### Data analysis for DDA purified virion samples

TimsTOF DDA files were searched in MSfragger using the LFQ-MBR workflow. Cysteine carbamylation was selected as fixed modification while N-term acetylation and deamidation were enabled as variable modification with a max of 3 variable modifications per peptide. Peptides of length 7 to 50 were searched again a database of phage, *Pseudomonas aeruginosa* plus contaminants. Decoys were generated by pseudo-inversion. Percolator was used for FDR-control at 1% PSM.

### Protein-protein interaction prediction from SEC-MS data

DIA-NN report were filtered at 1% library Q-value and, to infer protein quantities, the top2 peptides yielding the highest intra-protein correlation were averaged (sibling peptide correlation strategy). This step was performed across all samples to ensure the same peptides were used for every replicate and condition. The raw MS2 profiles were smoothed using a Savitzky-Golay filter and rescaled in a 0-1 range. A dot product matrix between all proteins was calculated and protein showing *r*^2^ *≥* 0.3 were selected as putative interactors for prediction. For every pair we calculated 5 features: (i) sliding window (q=6) correlation, (ii) fraction-wide intensity difference, (iii) peak shift, (iv) Euclidean distance and (v) contrast angle dot-product.

For prediction, we utilized a fully-connected neural network implemented in Tensorflow (https://www.tensorflow.org). Briefly, we set the input layer as number of features (147) followed by a fully connected layer with 100 neurons and a dropout layer (0.2 %) and a fully connected layer with 72 neurons. A final output layer using sigmoid as activation function was used for classifying coeluting and not-coeluting proteins. For training, a previously reported dataset was used[30]. To select for positive we utilized protein pairs in STRING using a combined score of 0.9 and experimental evidence, while negative were randomly selected. The DNN model was trained for 100 epochs using ADAM (learning rate = 0.001) and binary cross-entropy as loss function. Early stopping (patience = 20) to avoid overfitting. To further removed spuriously co-eluting PPIs after the prediction step, we calculated an equal number of decoy PPIs by randomly sampling the remaining proteins and utilized the DNN model to predict their coelution probability. We then utilized these two distributions to perform target-decoy competition (TDC) using posterior probabilities.

### ORFs prediction from nucleotide FASTA

EMBOSS v6.6.0.0 subroutine getorf was used to predict open reading frames (ORFs) with a minimum size of 50 AA. Existing annotated genes were removed from the predicted ORFs using bedtools subroutine subtract, allowing us to differentiate between existing and novel ORFs.

### Structural prediction and alignment for *ϕ*PA3 vRNAp

Protein complex prediction was performed using AlphaFold 2 (https://github. com/deepmind/alphafold). AF2 was run with full database size and the multimer preset. OpenMM energy minimization was performed to generate relaxed models and 5 models per complex were generated. Models were ranked by ipTM + TM and the PAE and LDDT were extracted for visualization. Each complex was submitted as a FASTA file, with proteins ordered from the longest to the shortest sequence. The alignment was performed using US-Align[49] (https://zhanggroup.org/US-align/) and the oligomer option was selected. Alignments of predicted complex structures (*ϕ*KZ vRNAp and 4 proteins *ϕ*PA3 vRNAp) were performed by multiple structure alignment (MSTA) using US-align with default parameters and a TM-cutoff of 0.45 was used to estimate topological similarities between the two structures. For visualization purposes, the structure of vRNAp (70GR) without PHIKZ123, which lacked homologs identification in *ϕ*PA3, was used as template in MatchMaker.

### Fluorescent microscopy of putative shell components

0.8% agar pads were supplemented with 0.5*µ*g/mL DAPI for phage DNA staining *P. aeruginosa* strain PAO1 expressing each of the fluorescent shell candidate constructs were grown in liquid culture supplemented with up to 0.05% arabinose (depending on optimal conditions for each construct) to induce construct expression until an OD of 0.5, and subsequently infected with *ϕ*KZ lysate for 50 min at 30°C before imaging. Microscopy was performed on an inverted epifluorescence (Ti2-E, Nikon, Tokyo, Japan) equipped with a Photometrics Prime 95B 25-mm camera and the Perfect Focus System (PFS). Images were acquired using Nikon Elements AR software (version 5.02.00). Cells were imaged through channels of phase contrast (200 ms exposure, for cell recognition), blue (DAPI, 50 ms exposure, for phage DNA), and green (GFP, 200 ms exposure, for mNeonGreen constructs) at 100x objective magnification. Final figure images were prepared in Fiji (version 2.1.0/1.53c)[63].

### Generation of *ϕ*KZ particles packaged with 3xFLAG fusions of *ϕ*KZ virion proteins

*ϕ*KZ particles packaged with virion proteins bearing a C-terminal 3xFLAG-tag were generated by adapting a protocol used to generate *ϕ*KZ particles packaged with mNeonGreen-tagged inner body proteins[18, 55]. PAO1 cells transformed with the appropriate pHERD30T*−*(PHIKZxxx)*−*3xFLAG construct were grown overnight in 3 mL LB supplemented with gentamicin (50 *µ*g/ml) at 37°C with aeration at 175 rpm. Cells were diluted 1:100 from a saturated overnight culture into 5mL LB supplemented with MgSO_4_ (10mM) and Gentamicin (50 *µ*g/ml) and grown for *≈* 2.5 hours at 37°C with aeration at 175 rpm. At OD600nm = 0.5-0.6 (3E8 CFU/mL), the bacterial cultures were infected with *ϕ*KZ (WT, MOI *≈* 1) for 2.5 hours. Thereafter 1 mL of chloroform was added to the cultures in a fume-hood and the cultures were incubated to with chloroform for 15 minutes (37°C, 175 rpm) The cell cultures were transferred to 15 mL falcon tubes and centrifuged at 6000xg for 15 min to pellet bacterial debris. The supernatant (containing bacteriophages in high titer) was carefully transferred to a fresh set of 50mL falcon tubes and centrifuged and 6000x*g* for 15 min to pellet any residual bacterial debris. Thereafter, 4 mL the supernatant was filtered and concentrated (*≈*10x) using Amicon-100 centrifugal filters to remove excess 3xFLAG-tagged proteins. The concentrated supernatant was used for western blot experiments.

### Western blot and blot analysis

PA01 cells were grown as previously described and upon reaching 0.5 OD (600 nm), gentamicin was added (50 *µ*g/ml) and the celles were chilled on ice for 5 minutes to stall translation. Thereafter PAO1 cells (*≈*1 OD equivalent) were infected with *ϕ*KZ particles packaged with virion proteins bearing a C-terminal 3xFLAG-tag (MOI *≈* 1) on ice for 10 minutes (to allow complete adsorption of virions onto cells) and then incubated at 30℃ for 15 minutes. Thereafter, the cell cultures were transferred to pre-chilled 15mL falcon tubes, centrifuged at 6000x*g*, 0°C for 5 minutes. The supernatant was discarded and the cell pellet was washed twice with 2 mL of pre-chilled (0°C) LB to remove excess unbound virions. The cell pellet was lysed in 100 *µ*L of lysis buffer (20 mM Tris, pH 7.5, 150 mM NaCl, 2% glycerol, 1% TTX-100, CompleteMini EDTA-free protease inhibitor cocktail). The lysed suspension was further sonicated on ice using a Q125 sonicator (10 pulses, 1s ON, 1s OFF, 30% amplitude). The cell lysate was centrifuged at 15000x*g* (15min, 0°C) to remove cellular debris. The clarified cellular lysate (100 *µ*L) was boiled with 33 *µ*LL of 4X Laemmli Buffer (with Beta-mercaptoethanol) for 10 min. 14 *µ*LL of lysate samples was loaded. For virion control samples, 10 *µ*LL of purified virions were boiled with 3.3 *µ*LL of 4X Laemmli Buffer (with Beta-mercaptoethanol) for 10 min and 2 *µ*LL of samples were loaded. SDS-PAGE gels were run with running buffer (100 mL 10X Tris-Glycine SDS Buffer, 900 mL Milli-Q water) at 130V for 1 hour (constant voltage setting). The SDS-PAGE gels were transferred onto 0.2 *µ*M PVDF membranes using a wet transfer (Transfer Buffer: 100 mL 10X Tris-Glycine Buffer, 200 mL methanol, 700 mL Milli-Q water; 100V, 1 hour, 4°C). The membranes were incubated with blocking buffer (5% Omniblock milk, non fat-dry in 1X TBST (200 mL Tris Buffer Saline, 0.20 mL Tween-20)) for 1 hour at room temperature. Thereafter the blocking buffer was discarded and the membranes were incubated with 1:1000 dilutions of mouse anti-FLAG M2 antibody (Sigma-Aldrich) in 1X TBST (overnight, 4°C, with constant shaking). Thereafter the membranes were washed thrice for 10 min with TBST and incubated with 1:3000 dilution of Goat anti-mouse HRP (Thermo Fischer) in blocking buffer for 1 hour at room temperature with constant shaking. Finally, the membranes were washed thrice for 10 min with TBST and incubated with Clarity Western ECL substrate. Membranes were imaged on an Azure 500 imager.

